# Efficient Deep Learning Models for Predicting Individualized Task Activation from Resting-State Functional Connectivity

**DOI:** 10.1101/2024.09.10.612309

**Authors:** Soren J. Madsen, Young-Eun Lee, Shaun K. L. Quah, Lucina Q. Uddin, Jeanette A. Mumford, Deanna M. Barch, Damien A. Fair, Ian H. Gotlib, Russell A. Poldrack, Amy Kuceyeski, Manish Saggar

**Author notes:** The authors contributed equally to this work (Soren J. Madsen, Young-Eun Lee).

## Abstract

Deep learning models have demonstrated the potential to predict task-evoked brain activation from resting-state fMRI, offering a pathway toward individualized brain mapping without requiring task-based data. In this study, we systematically evaluate architectural strategies for improving the efficiency and scalability of such models. Using data from the Human Connectome Project, we replicate the BrainSurfCNN framework and introduce two extensions: BrainSERF, which incorporates channel-wise attention through squeeze-and-excitation modules, and BrainSurfGCN, a graph-based model that leverages cortical mesh topology for efficient message passing. Across multiple evaluation metrics, including spatial correlation, Dice score, Dice AUC, and subject identification accuracy, all models achieve comparable predictive performance. Despite similar accuracy, the proposed models offer distinct advantages. BrainSERF provides modest improvements in capturing individual-specific features, while BrainSurfGCN achieves substantial reductions in model size and training time, highlighting a favorable trade-off between performance and computational efficiency. Beyond architectural comparisons, we investigate factors driving variability in prediction accuracy. We find that behavioral task performance, resting-state data quality, and inter-subject variability in task activation jointly constrain prediction fidelity. In particular, contrasts with lower signal reliability and higher variability exhibit reduced predictability across all models. Together, these findings demonstrate that incorporating topological and functional structural priors can improve the efficiency of deep learning models without sacrificing accuracy, while also emphasizing that prediction performance is fundamentally limited by the reliability of the underlying neural signals.

## 1 Introduction

One of the foundational questions in cognitive neuroscience is whether the brain’s intrinsic organization, observable during rest, encodes information predictive of how that same brain will respond during active cognitive tasks. If intrinsic functional connectivity reflects stable traits that constrain task-evoked activation, then models trained on resting-state data should be able to accurately predict task contrast maps at the individual level (Cole et al., 2016; Ngo et al., 2022; Osher et al., 2016; Tavor et al., 2016). This hypothesis has significant implications for understanding brain-behavior relationships, characterizing individual differences, and developing precision neuroimaging tools, particularly for populations that are unable to perform scanner-based tasks. At the same time, validating such models provides a data-driven approach to test assumptions about the relationship between intrinsic and task-evoked activity.

Prior work has demonstrated that the spatial organization of resting-state networks closely mirrors the brain’s task-evoked architecture (Smith et al., 2009; Smith-Collins et al., 2015; Sui et al., 2009), suggesting that intrinsic functional connectivity may serve as a latent scaffold for cognitive function. This raises the possibility that resting-state functional magnetic resonance imaging (rsfMRI) contains sufficient information to reconstruct task activation maps at the individual level, effectively using the brain at rest to infer how it behaves in action. In addition to its practical utility in cases where task data are unavailable or infeasible (e.g., clinical or pediatric populations), such predictions offer a powerful test of the stability, specificity, and behavioral relevance of intrinsic connectivity (Chow et al., 2025; Hearne et al., 2021; R. Jiang et al., 2020; Savage et al., 2024; Serin et al., 2025). Recent reviews have further summarized advances in this area and highlighted its expanding methodological scope and applications (Bernstein-Eliav and Tavor, 2024), extending connectome-based prediction to include mechanistic frameworks and deep learning approaches. The feasibility of this approach has also been demonstrated in clinical populations, including pre-surgical mapping and psychiatric cohorts (Jones et al., 2017; Tik et al., 2021), highlighting its potential translational utility.

This line of inquiry also has significant implications for translational research frameworks such as the NIMH Research Domain Criteria (RDoC), which seek to redefine mental illness in terms of dimensional, neurobiologically grounded constructs (Cuthbert and Insel, 2013; Insel, 2014). Task-fMRI has traditionally played a central role in identifying brain circuits linked to RDoC domains, yet no single individual completes all relevant tasks (Quah, Madsen, et al., 2025; Quah, Jo, et al., 2025). Accurate rest-to-task prediction could enable virtual approximation of these activation patterns, potentially allowing comprehensive, individualized neural profiling without requiring data acquisition across all RDoC task domains. Thus, predictive models offer both a mechanistic window into brain organization and a scalable tool for next-generation clinical neuroscience.

Foundational work by Tavor et al., 2016 and Cole et al., 2016 demonstrated that task activation maps could be predicted from resting-state functional connectivity using linear models, establishing a baseline for rest-to-task inference. These studies showed that intrinsic connectivity patterns contain behaviorally relevant information. Building on this foundation, Ngo et al., 2022 introduced BrainSurfCNN, a deep learning model that integrates U-Net-style architecture with surface-based mesh convolutions to significantly improve individual-level prediction accuracy. While technically impressive, such models raise two central questions that remain unresolved: (1) Can architectural changes enhance the fidelity or efficiency of these predictions? And (2) why are some individuals or task contrasts more predictable than others?

To address the first question, we examine whether these architectural changes can enhance the fidelity or efficiency of rest-to-task predictions. We introduce two new models built upon distinct principles. The first, BrainSERF, augments BrainSurfCNN with squeeze-and-excitation (SE) modules (Hu et al., 2018), which implement dynamic channel-wise feature reweighting. This mechanism performs adaptive channel-wise feature reweighting that modulate information flow based on contextual salience. The second model, BrainSurfGCN, utilizes graph convolution (Kipf and Welling, 2016) and models information propagation over the cortical mesh using graph-based operations by treating the cortical surface as a graph and explicitly modeling localized, neighborhood-based information flow, thereby explicitly modeling localized, neighborhood-based information flow on the cortical mesh.

To address the second question, we investigate why rest-to-task prediction succeeds for some individuals or task contrasts but not others. Specifically, we examine whether variability in model performance may reflect meaningful differences in cognitive engagement, neural signal reliability, and the degree to which intrinsic connectivity scaffolds task-evoked activity. To address this, we evaluate how behavioral performance and scan quality influence prediction accuracy, and we examine the cortical distribution of prediction errors in relation to known anatomical noise profiles. These analyses aim to reveal when and where brain dynamics and individual traits facilitate or constrain successful decoding of task-related activity from resting-state data.

## 2 Method

### 2.1 Dataset and Preprocessing

We used de-identified, publically available data from the HCP dataset (Van Essen et al., 2013) to train our models in the same way as BrainSurfCNN. The study was approved by the Stanford University Institutional Review Board.

Specifically, as input to our models, we used the FIX-cleaned, 3T rsfMRI data acquired in four 14.4-minute runs, each with 1200 time points per session per subject. The acquisition and preprocessing methods for the HCP dataset have been described elsewhere in (Barch et al., 2013; Glasser et al., 2013; Smith et al., 2013), where rsfMRI preprocessing includes highpass filtering, MELODIC ICA, and FIX for component classification. We employ the data augmentation scheme as proposed in (Ngo et al., 2022), in which samples of the resting state data are drawn from all four recordings of resting state data. This aggregation of recording data amounts to 4,800 time points split into eight samples of 600 time points from which the functional connectivity is determined. These eight samples allow us to extend the training set of our data by uniformly sampling these eight chunks in each training epoch.

Group-level parcellations derived from spatial ICA were also released by the HCP, and we examined the performance of our models trained on various component parcellations for computing the functional connectomes. We create new training datasets derived from 15, 25, 50, and 100 independent components using these spatial components. As HCP’s tfMRI data spans seven task domains, and, following Tavor et al., 2016 and Ngo et al., 2022, we obtained 47 unique contrasts after excluding redundant task contrasts. This approach allowed us to explore the impact of independent components on the model’s ability to predict task contrasts accurately on both an individual and a group level of analysis.

Additionally, we only included subjects with all four resting-state runs and all 47 task contrasts, which gave us 919 subjects for training set, of which five subjects were used for a validation set (Ngo et al., 2022). The use of the validation set prevents overfitting our models during the training process.

Following the filtering approach described by Ngo et al., 2022), our test-retest sample consisted of 39 out of 46 available subjects who met the data completeness criteria, specifically having all four rsfMRI runs and 47 tfMRI contrasts. We refer to the contrasts from the first visit as the “test set” and those from the second visit as the “repeat dataset,” adopting BrainSurfCNN terminology. The test set participants do not overlap with those in our training set. We used the repeat dataset as an optimal predictor of groundtruth task activation (i.e., non-model-based or model-free). A contrast taken from the same subject in a different session represents the best possible model for individual variability, providing an approximate noise ceiling, particularly at the group level, although individual-level comparisons may deviate due to measurement variability. Notably, resting-state data from the repeat sessions were not used as model input, but were instead used as a reference to estimate the noise ceiling. In addition, we included a group-average baseline as a non-individualized reference. Specifically, for each task contrast, we computed the average activation map across training subjects and used this as a prediction for each individual subject. This baseline reflects population-level activation patterns while removing subject-specific variability.

In the proposed deep learning framework, the input data are structured as multi-channel fsLR polyhedral meshes, each consisting of 32,492 vertices, to accommodate the high-dimensional spatial topology of cerebral cortex representations. Here, “fsLR” denotes FreeSurfer-derived cortical surface meshes aligned to a standardized template (fsLR template) across the left and right hemispheres. Each channel within these meshes is dedicated to an independent component derived from functional MRI data preprocessing, serving as a distinct feature for the model’s input layer. For instance, in a scenario where the model is configured to analyze 50 independent components, the total input dimensionality would amount to 100 channels, reflecting the bilateral nature of cerebral anatomy with a separate mesh for each hemisphere. The model’s output mirrors the input’s structural format, generating a polyhedral mesh with 32,492 vertices. However, in the output mesh, each vertex’s value across the channels indicates the predicted task contrast for the corresponding location in the brain hemisphere. Given that the study focuses on predicting 47 distinct task contrasts, the model’s output dimensionality is adjusted to feature 94 channels, with each pair of channels representing the predicted contrasts for the left and right hemispheres, respectively.

### 2.2 Models

We evaluated three models, BrainSurfCNN, BrainSERF, and BrainSurfGCN, for predicting task contrast maps. These models differ in how they represent and propagate spatial information on the cortical surface. BrainSurfCNN models local spatial structure via mesh convolutions. BrainSERF extends this framework by enhancing feature selectivity through channel-wise recalibration. BrainSurfGCN, in contrast, propagates information across the cortical graph using message passing. We describe each model in detail below.

#### 2.2.1 BrainSurfCNN

BrainSurfCNN is a mesh-based encoder-decoder convolutional network that maps resting-state IC representations to task contrast maps using hierarchical surface convolutions. The development of BrainSurfCNN integrates the foundational principles of a U-Net architecture (Milletari et al., 2016; Ronneberger et al., 2015), a prevalent framework in medical image segmentation, with advanced mesh convolution techniques to facilitate the analysis of brain surface data. This innovative approach is significantly influenced by the pioneering work on convolution for spherical meshes called UGSCNN, as detailed by C. Jiang et al., 2019.

The architecture adopts a hierarchical encoder-decoder structure composed of downsampling and up-sampling pathways, which leverage mesh convolutional operations to effectively capture spatial features across multiple resolutions. Importantly, it incorporates a Residual Pooling Block (implemented as ResPoolBlock), which combines residual connections, mesh convolutions, and pooling operations to enhance robust multi-scale feature learning. The primary objective of BrainSurfCNN (Ngo et al., 2022) is to leverage the spatial hierarchies inherent in polyhedral meshes to predict task-related activation patterns, represented as contrast maps, from resting-state IC maps.

In our study, we replicated the BrainSurfCNN model using the source code made publicly available by the original authors on GitHub ^1^. This replication process was not merely an exercise in model reconstruction but a deliberate effort to validate and extend the model’s application. A cornerstone of our analysis involved a systematic examination of BrainSurfCNN’s performance across a spectrum of resting-state scans characterized by varying numbers of independent components.

#### 2.2.2 BrainSERF

BrainSERF extends BrainSurfCNN by incorporating channel-wise feature recalibration through squeeze-and-excitation (SE) modules. The SE attention mechanism has emerged as a pivotal innovation in the field of computer vision, significantly enhancing the representational power of convolutional neural networks by enabling channel-wise feature recalibration (Hu et al., 2018). This technique introduces trainable scaling coefficients that adaptively adjust the weighting of each channel in the network’s feature maps, thereby allowing the model to emphasize informative features and suppress less useful ones dynamically. The SE attention mechanism operates by first ‘squeezing’ global spatial information into a channel descriptor through global average pooling, followed by ‘excitation’ operations, fully connected layers that capture channel-wise dependencies. Given two weight matrices *W*_*sq*_ and *W*_*ex*_, we perform the squeeze operation by globally pooling each mesh so that our representation of input *X* transforms from shape ℝ^32,492x*C*^ to ℝ^1x*C*^ where *C* represents the number of hidden channels. Next, we define a ratio *r* such that the matrix *W*_*sq*_ transforms the data from ℝ^1x*C*^ to 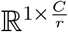. We finish the ‘squeezing’ with a Tanh activation to capture negative values to get X’. To perform the ‘excitation’ step, we multiply the transformed X’ by *W*_*ex*_ to expand the channel-wise axis back to a matrix in ℝ^1x*C*^ to get *S*. We applied a Tanh activation function again to capture negative values. This matrix S represents a scaling on the channel-wise axis. We channel-wise multiply X ∈ ℝ^32>492x*C*^ by S to achieve the output *X*_*scaled*_, a channel-wise rescaled form of the input X. The intuition with this mechanism is to periodically rescale the hidden representation of the mesh channels and highlight or suppress entire latent activation patterns.

These operations produce scaling coefficients applied to the original feature maps, effectively allowing the network to perform self-attention across channels. In our network architecture, we implement SE attention before the mesh convolutional layers during the coarsening stage of the mesh, inspired by the work in Wang et al., 2021. The network additionally incorporates the Residual Pooling Block, as described in BrainSurfCNN, to further enhance hierarchical feature learning through residual connections and mesh-based pooling.

Furthermore, we have innovated beyond traditional activation functions by transitioning from Rectified Linear Unit (ReLU) activations to Hyperbolic Tangent (Tanh) activations throughout our network. This adjustment is primarily motivated by the need to accommodate negative values within the SE attention mechanism and the predicted task contrasts. The Tanh activation function enables our network to capture a broader spectrum of feature dynamics, including both positive and negative activations that are present in the task contrast maps. The BrainSERF architecture is described in Figure |1a, and additional training hyperparameters, command-line arguments, and example usage scripts are provided on our GitHub repository: https://github.com/braindynamicslab/dl-task-contrast-prediction

#### 2.2.3 BrainSurfGCN

BrainSurfGCN is a graph convolutional network that operates directly on the cortical mesh, using message passing to propagate information across neighboring vertices.

In predicting task contrast activation maps from rsfMRI, employing graph neural networks offers several compelling advantages that align with the intrinsic properties of brain data and the objectives of neuroimaging analysis (Zhang et al., 2021). The human brain can be conceptualized as a complex network, with nodes representing different brain regions and edges representing functional or structural connections between these regions. This graph-centric view is inherently compatible with the structure of GNNs. We define a graph below.

##### Definition 1

*An undirected graph G can be defined as G =* (𝒱, ℰ), *where v* ∈ 𝒱 *represents a node with feature size in* ℝ^*C*^ *and e* ∈ ℰ *represents an undirected edge between two vertices* v *and v*_*j*_.

We leverage the spherical mesh used in Ngo et al., 2022 as the graph’s structure for each input, such that | 𝒱 | = 32,492 mesh. Thus, our input data has two parts. The node-wise representation, 𝒱, is represented as a matrix in ℝ^| 𝒱 |*x* 𝒸^, where each node *v* ∈ 𝒱 has *C* learnable parameters. The edge-wise representation is a list of tuples of vertices that are connected such that edge e ∈ ℰ connecting vertices v_*i*_ and *V*_*j*_ is represented as *(i,j*), and ℰ ∈ ℤ^2x| ℰ |^.

We build BrainSurfGCN upon the work of Kipf and Welling, 2016. The architecture we use for this study builds upon the implementation of the Graph Convolution Layer seen in PyTorch Geometric (Fey and Lenssen, 2019), and we build a network composed of what we call a BDLayer (described in Figure S1). The model consists of multiple graph-based layers, each composed of a GCN Layer, a LeakyReLU activation, and a BatchNorm layer. The BatchNorm layer accelerates learning by reducing internal covariate shift (Ioffe and Szegedy, 2015) through the recentering of features using the formula:

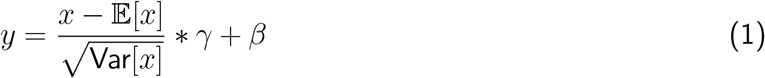

where *γ* and *β* are learnable parameters, this layer acts along a node’s feature-wise axis. The GCN Layer defines the update of a node’s feature space from *X* to *X′* as

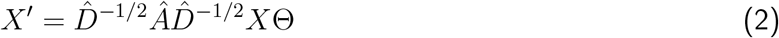

where *Â = A* + *I* denotes an adjacency matrix with added self-loops, 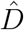 denotes the diagonal degree matrix of *Â*, and e denotes the learnable weight matrix. The adjacency matrix *A* is formed such that *A*_*ij*_ = 1 if *e*_*ij*_ ∈ ℰ, and A_*j*_ = 0 if *e*_*i j*_ ∉ *ℰ*. This operation allows us to pass information between vertices on the mesh at a distance of 1 hop per layer. To ensure that node information is disseminated across the entire graph, we use 8 BDLayers. Additionally, we believe that spatial information is important in predicting the activation on the mesh, so we included the XYZ coordinates of each vertex as 3 additional channels for our input data. These coordinates represent the locations of the vertices of the polyhedral sphere centered on the origin (0,0,0) with a radius of 1. Work in Liu et al., 2018 also suggests that including spatial information can improve convolution operations. With a resting-state map of 50 ICs per hemisphere, our input feature space lies in ℝ^32>492x(2*50+3)^. Figure 1b shows our architecture from a high-level perspective, complete with the input and output feature spaces.

**Figure 1:**
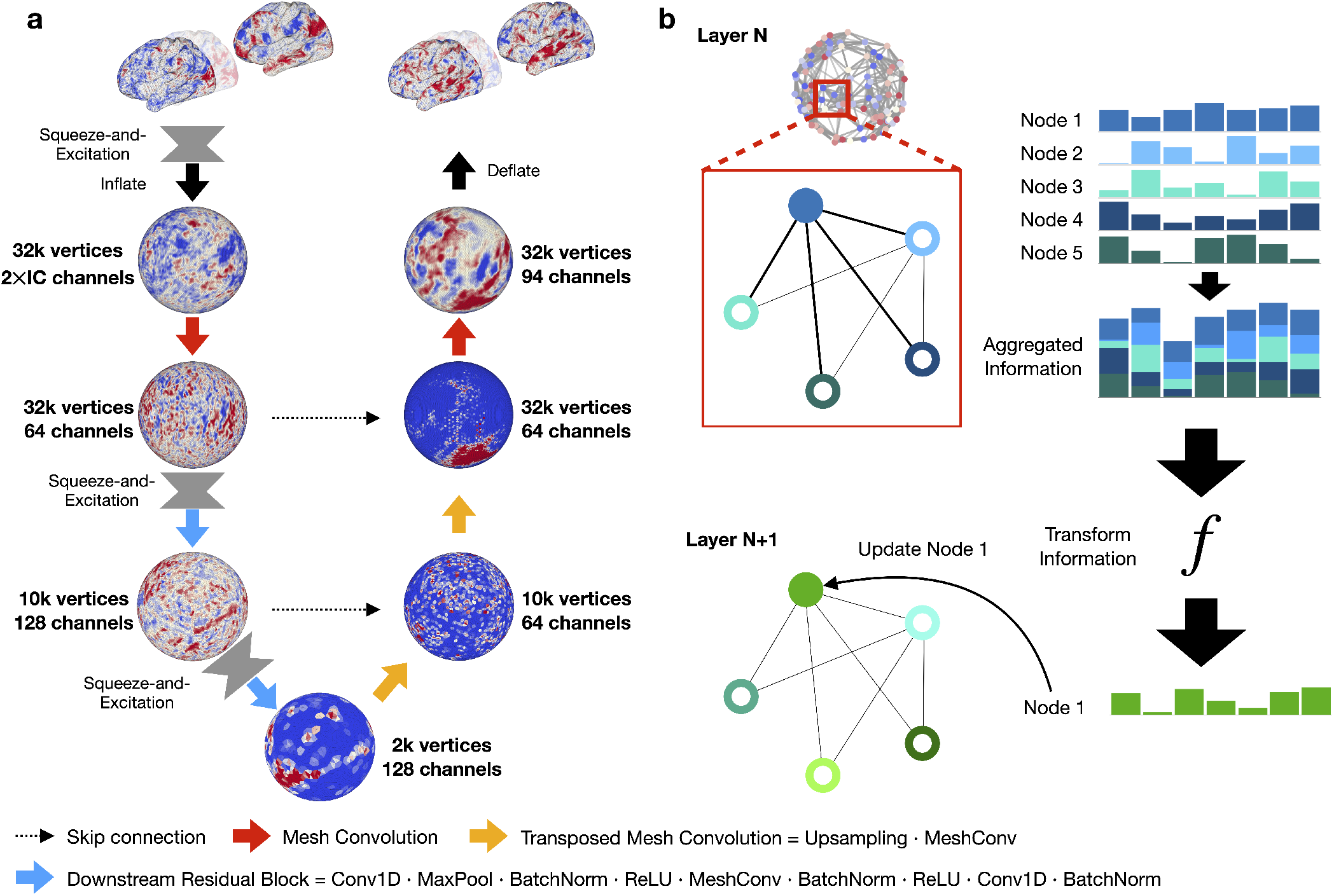
Overview of BrainSERF and BrainSurfGCN architecture for cortical mesh-based fMRI analysis. a. In the BrainSERF model, input surface-based fMRI data are processed through a hierarchical mesh convolutional network, incorporating SE modules and skip connections. Vertices and channel dimensions are progressively transformed using mesh convolution, residual blocks, and upsampling operations. b. In each BrainSurfGCN layer, node features in the cortical mesh are aggregated, transformed, and updated according to graph connectivity. Aggregated node information is passed through nonlinear transformations to generate updated node representations for subsequent layers. This framework enables multi-scale integration and hierarchical feature extraction from full-brain functional mesh data.

### 2.3 Training Details

All the models trained in these experiments used the same set of hyperparameters and training regimes. Models were developed and trained in PyTorch (Paszke et al., 2019), and we used Adam (Kingma and Ba, 2014) for optimization. We utilized a learning rate of 0.001 and trained the models for 50 epochs using MSE loss to promote task contrast reconstruction. We used the model with the best-predicted correlation on the validation set at the end of 50 epochs for evaluation.

After performing some preliminary sanity checks (e.g., correlation between prediction and ground truth), we continued with the next phase of training. In this second step, we used the trained model to compute and average the MSE across the training set for same-subject reconstruction loss, α, and across-subject reconstruction loss, *γ*, to be used as hyperparameters in the next step of training that used a reconstructive-contrastive (RC) loss function proposed by Ngo et al., 2022. Given a mini-batch of *N* samples, 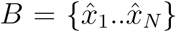, in which 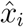 is the target multi-channel contrast image of subject *i*, the RC loss function is defined as

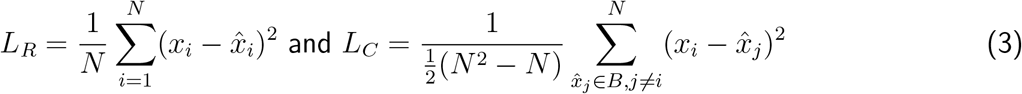

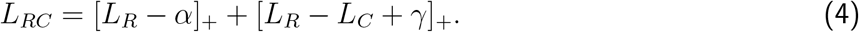

Using the RC loss described in Equation 4, we train the model again for 50 epochs. Before training, we initialize values for *𝒱* and *γ* from the losses computed over the training set. These values are updated every 10 epochs such that:

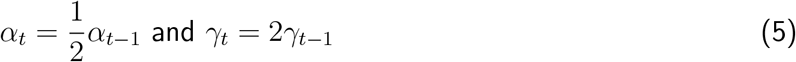

until *α* reaches a minimum of 1 and *γ* reaches a maximum of 10. Using this RC loss function, we encouraged the model to differentiate task contrast reconstructions between subjects, thereby capturing individual differences.

### 2.4 Evaluation

To evaluate the efficacy of our model in predicting task contrast maps from resting-state fMRI data, we employ three distinct metrics: spatial correlation, Dice coefficient (Dice score; also referred to as F1 score), Dice AUC score for all thresholds, and subject identification accuracy. These metrics offer a comprehensive assessment of the model’s performance, each addressing different aspects of prediction quality.

The spatial correlation metric primarily measures the linear relationship between the model’s predicted task contrasts and the empirically observed (ground truth) contrasts. This metric was chosen for replication to compare the results from Ngo et al., 2022. By computing Pearson’s correlation coefficient, we quantify how the predicted values co-vary with the actual task contrasts across the mesh’s vertices. A high correlation coefficient indicates that the model effectively captures the spatial distribution of neural activations associated with specific tasks, mirroring the patterns observed in the ground truth data. A crucial aspect of the performance is ensuring that within-subject covariance is more similar than inter-subject covariance.

The Dice coefficient (Dice score) enhances our evaluation by measuring spatial overlap between the predicted and ground truth task contrast activation patterns. To calculate this metric, we first apply a series of thresholds to the activation values at each vertex, classifying them as ‘activated’ or ‘non-activated.’ For each threshold, the Dice score is computed as the harmonic mean of precision and recall between the two binary sets of activated vertices. Thus, the specificity of predicted activation on the contrasts is computed over a range of threshold values, ensuring predicted regions with higher activation (from a higher threshold) overlap with the higher activations in the ground truth contrasts. Thereby, we capture another measurement, in addition to correlation, related to both the spatial distribution and the relative magnitude of the cortical activation.

To obtain a threshold-independent summary, we further compute the Dice AUC by integrating Dice scores across all evaluated thresholds. This provides a more robust assessment of spatial overlap that does not depend on a single threshold choice, while still capturing both the spatial distribution and relative magnitude of cortical activation.

To assess the model’s capability in capturing individual-specific features within the task contrasts, we use an evaluation method introduced by Ngo et al., 2022 based on subject identification accuracy. This approach involves computing the correlation between each predicted task contrast and all available ground truth contrasts across subjects for the same task. The identification of the subject is deemed correct if the highest correlation corresponds to the correct subject’s ground truth contrast compared to those for other subjects. This metric reflects the model’s sensitivity to individual differences in brain activation patterns, highlighting its potential for personalized neuroimaging analysis. Moreover, we computed this accuracy for each quantity of ICs used in training.

## 3 Results

In this section, we address two core questions central to our investigation: (1) Can architectural changes improve the fidelity and efficiency of predicting task-evoked brain activation from resting-state connectivity? and (2) Why does prediction performance vary across individuals and cognitive tasks?

To this end, we organize our results in a three-part sequence: (i) Architectural Design and Trade-offs: We first evaluate predictive performance across three model architectures, including BrainSurfCNN (baseline), BrainSERF, and BrainSurfGCN, highlighting efficiency improvements. Model predictions are compared against both a repeat scan from the same subject, serving as a noise ceiling, and a group-average baseline, serving as a non-individualized reference. (ii) Model Evaluation: We then assess subject-level prediction specificity through a subject identification analysis, offering a complementary measure of how well each model captures individualized activation patterns. (iii) Mechanisms of Variability: Finally, we examine factors that drive individual and contrast-level differences in prediction performance, including task engagement, data quality, and inter-subject variability in activation patterns. All results were computed using 50 ICs for consistency.

### 3.1 Architectural Design and Trade-offs

We first evaluated how well each model, including BrainSurfCNN, BrainSERF, and BrainSurfGCN, could predict individual-level task activation maps from resting-state connectivity patterns. As shown in Figure 2 for one representative participant, all three models qualitatively captured key spatial features of task-evoked responses, such as somatomotor activation during the Motor task and activation in association cortices during the Social task. BrainSurfGCN achieved comparable spatial localization and alignment with ground truth, while substantially reducing model complexity and computational cost. The thresholded task activation maps for the other IC settings (e.g., 15, 25, and 100) across all predictions for the seven task contrasts are presented in the Supplementary Materials (Figures S2, S3, and S4). Performance was not substantially affected by the choice of IC dimensionality.

**Figure 2:**
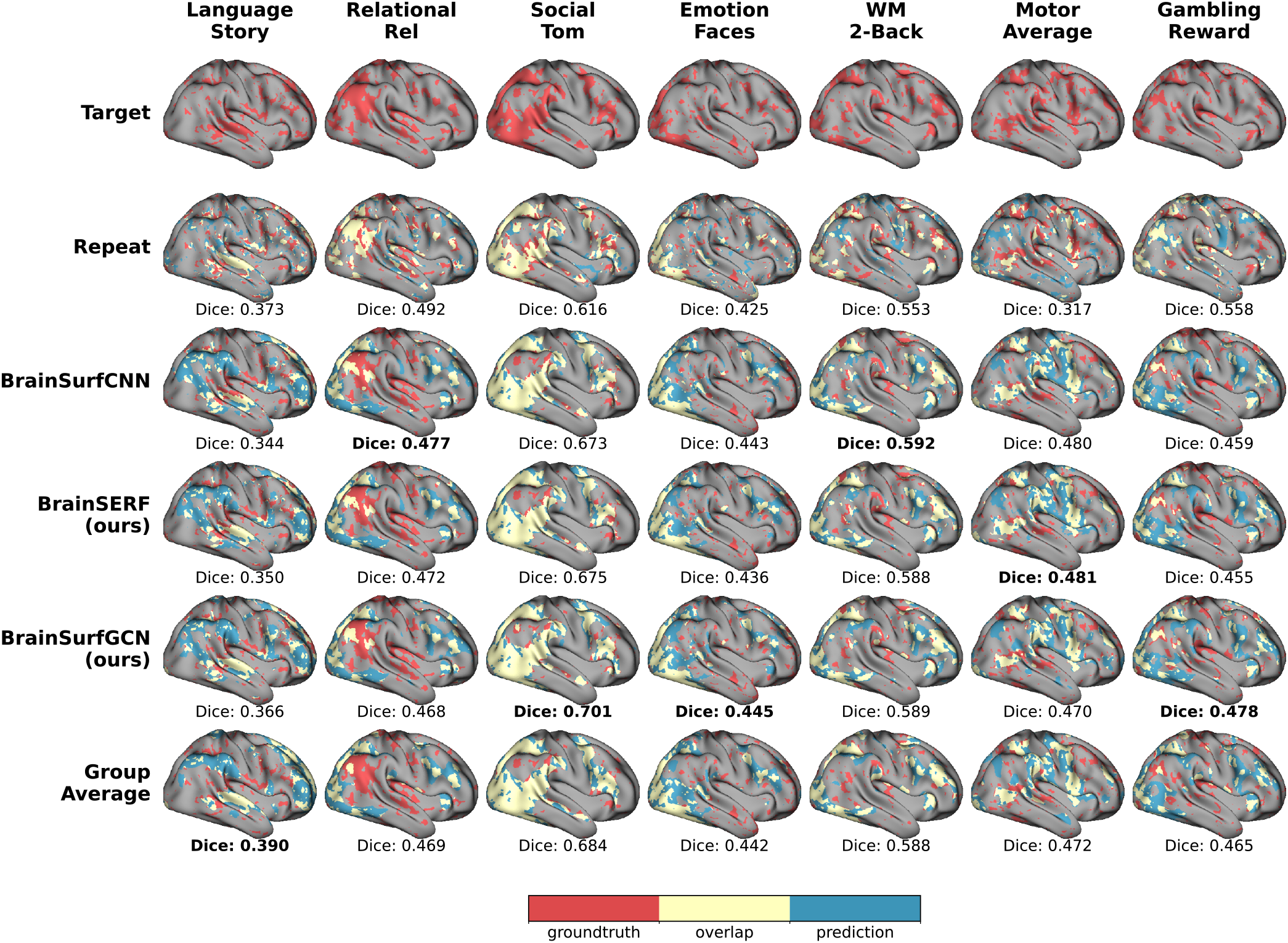
Thresholded task activation maps for 7 task contrasts using threshold of 25%. We compare ground truth, repeat, group average, and models’ predicted activation for an individual (subject 917255). The thresholded activation maps on the right hemisphere (lateral view) represent, respectively, Language: Story, Relational Processing: Relational, Social Cognition: Theory of Mind, Emotion: Faces, Working Memory: 2-Back, Motor: Average, and Gambling: Reward. Ground truth (target), repeat scans, group average, and predictions from BrainSurfCNN, BrainSERF, and BrainSurfGCN are presented with colors indicating ground truth, model prediction, and overlap. Dice scores are displayed beneath each row, quantifying prediction fidelity across thresholds. The highest Dice value within each column is bolded to highlight the best-performing models.

To quantitatively evaluate model performance, we computed Dice score between predicted and ground-truth activation maps across multiple thresholds (5%-50%, in 5% increments), following the procedure established in Ngo et al., 2022. Thresholding allows us to isolate task-relevant signal while mitigating the influence of noise. Figure 3 shows example outputs for three thresholds (10%, 25%, 50%) from the Social Cognition: Theory of Mind contrast, illustrating how prediction sensitivity changes with threshold level.

**Figure 3:**
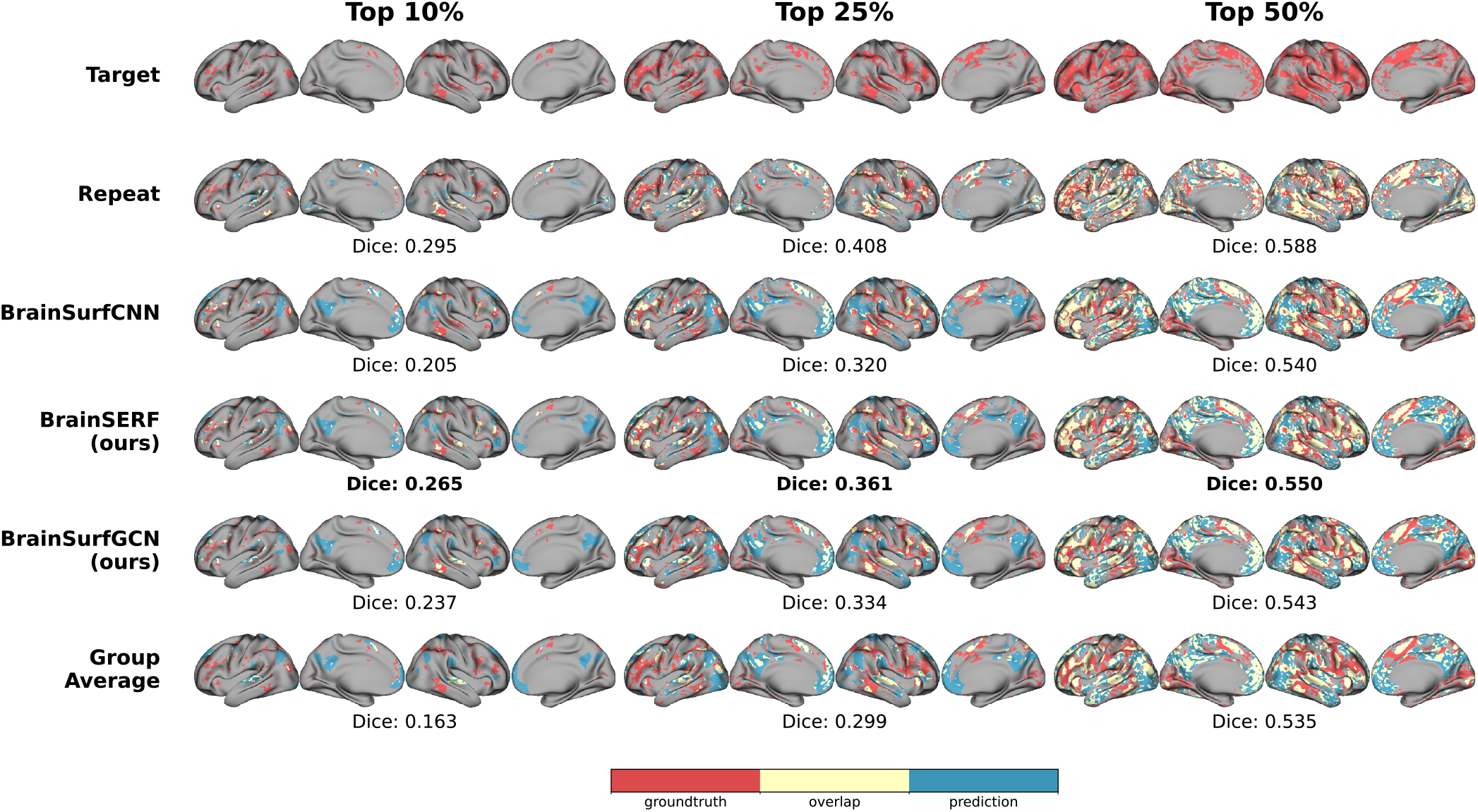
Thresholded task activation maps for Social Cognition: Theory of Mind. Activation maps for an individual (subject 917255) are thresholded at three levels (top 10%, top 25%, and top 50%) to illustrate how effectively each model captures the spatial distribution of task-evoked responses. Ground truth (target), repeat scans, group average, and predictions from BrainSurfCNN, BrainSERF, and BrainSurfGCN are shown with colors indicating ground truth, model prediction, and overlap. Dice scores are displayed beneath each row, quantifying prediction fidelity across thresholds. The highest Dice value within each column is bolded to highlight the best-performing models.

We computed Dice scores across thresholds from 5% to 50% in 5 percent steps to characterize how prediction quality changes with varying activation selectivity. As shown in Figure 4, these curves provide a clearer view of model behavior across activation intensities. Across different task contrasts, we observe three representative patterns: (1) convergence between repeat and group average performance, (2) comparable performance between model predictions and repeat, and (3) improved performance of model predictions over both repeat and group average. In several contrasts, model predictions slightly exceeded the repeat dataset, which represents the empirical noise ceiling. This pattern, also noted by Ngo et al., 2022, likely arises because repeat scans contain session-to-session variability and small spatial shifts in activation, lowering their measurable overlap. The smoother and less noisy nature of model predictions can therefore yield higher Dice scores without violating theoretical limits. All Dice scores for every final model across all task contrasts are provided in the Supplementary Figure S6.

**Figure 4:**
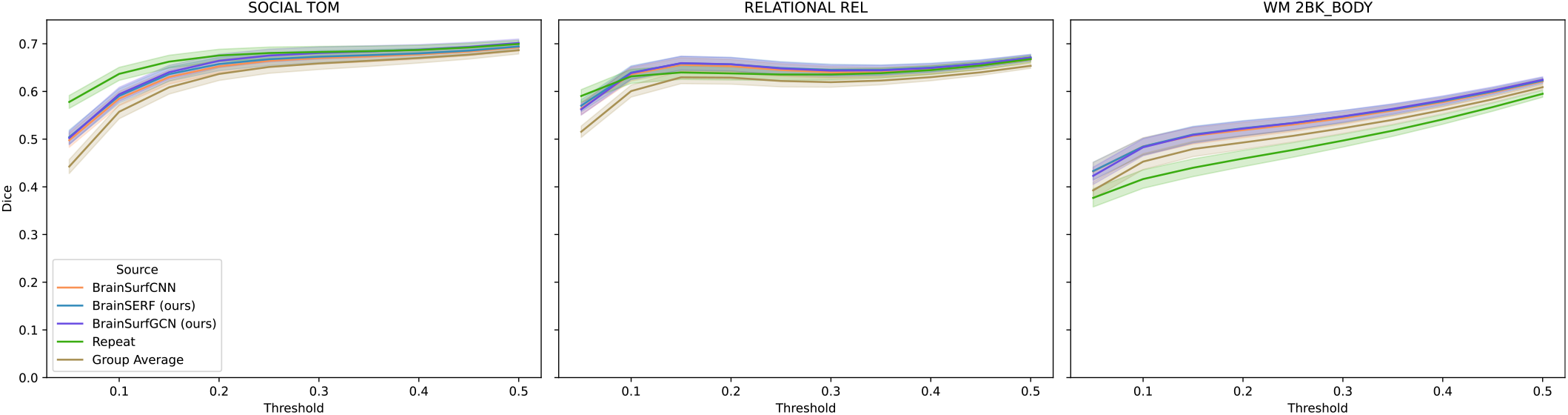
Dice score plots for three selected tasks showing variations in performance across task contrasts. Each plot compares model predictions with the repeat condition and group average baseline across thresholds from 5% to 50% for Social Cognition: Theory of Mind, Relational: Relational, and Working Memory: 2-Back Body. The shaded regions denote standard deviations in the dice scores across test subjects.

To summarize performance across thresholds, we calculated the area under the Dice curve (Dice AUC), with a maximum possible value of 0.45 based on perfect overlap across the integration range. These values are shown in Figure 5 and provide a concise metric for comparing model performance across tasks and individuals. We also examined unthresholded predictions for baseline reference.

**Figure 5:**
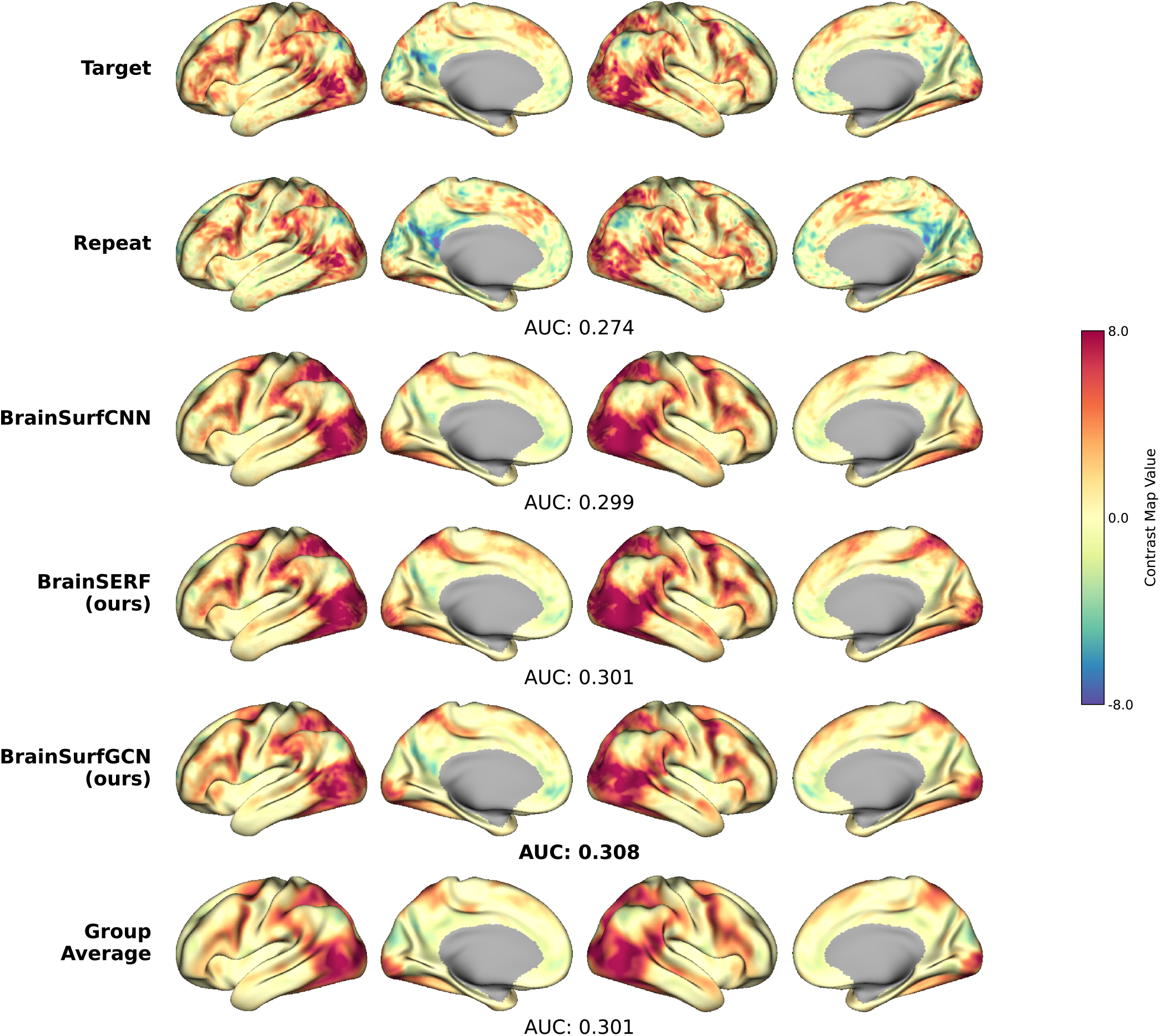
Unthresholded task activation for Social Cognition: Theory of Mind. We compare ground truth, repeat, group average, and model-predicted activation for an individual (subject 917255). Model predictions are much smoother on the whole than the ground truth or repeat activations. Though smoother, the models perform higher in the Dice AUC than the repeat contrasts. All activation maps are displayed in z-score units.

Finally, extended results, including per-task Dice AUC and additional visualizations, are available in Supplementary Tables S1 and S2 and Figure S7-S15, providing a more granular view of how each model performs across contrasts and individuals.

Finally, we quantified the computational footprint of each model to highlight trade-offs between performance and efficiency. Table 1 reports the number of trainable parameters across architectures and ICs. Notably, BrainSurfGCN achieved substantial reductions in parameters. This is attributable to its mesh-based design, which allows parameter sharing across nodes and avoids the need to store full mesh gradients. Importantly, model size remained largely stable across different IC configurations, since operations scale with fixed hidden-state sizes rather than the number of ICs directly.

**Table 1:**
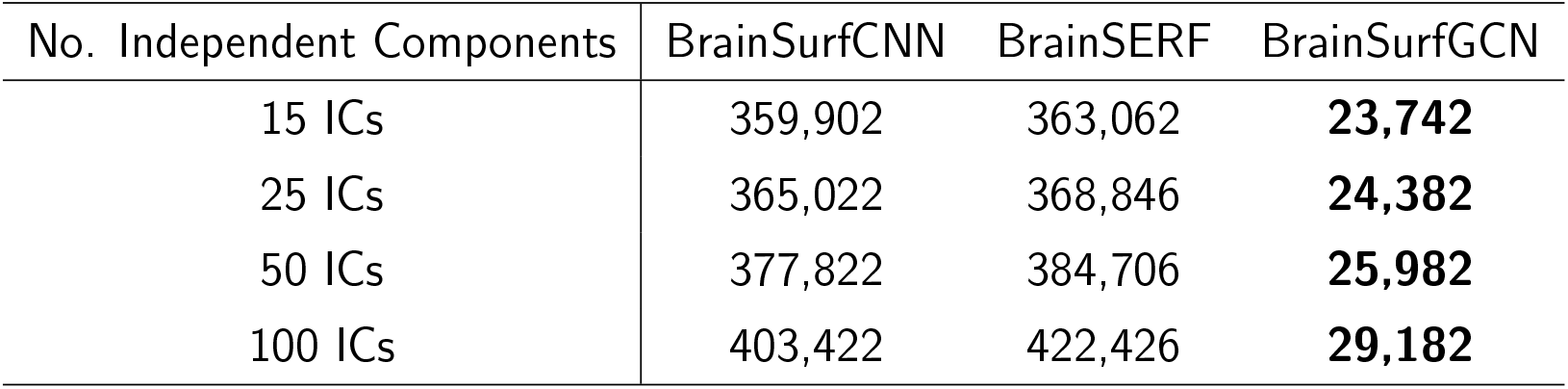
Number of Trainable Parameters for Each Model.

| No. Independent Components | BrainSurfCNN | BrainSERF | BrainSurfGCN |
| --- | --- | --- | --- |
| 15 ICs | 359,902 | 363,062 | **23,742** |
| 25 ICs | 365,022 | 368,846 | **24,382** |
| 50 ICs | 377,822 | 384,706 | **25,982** |
| 100 ICs | 403,422 | 422,426 | **29,182** |

### 3.2 Evaluation of Predictive Fidelity - Subject-Level Specificity

To evaluate the individual specificity of model predictions, we examined both the spatial correlation between predicted and ground-truth task activation maps and the subject identification accuracy. This analysis assesses whether the models capture unique, individualized features of brain function that distinguish one subject from another.

We first computed the spatial correlation difference: for each predicted contrast, we compared its correlation with the corresponding ground-truth contrast from the same subject versus those from different subjects. A high correlation difference indicates that the model more accurately captures idiosyncratic features of the target individual. As shown in Figure 6, all three models exhibit strong within-subject correlation relative to others, suggesting that they reliably capture individually localized patterns of task-evoked activity.

**Figure 6:**
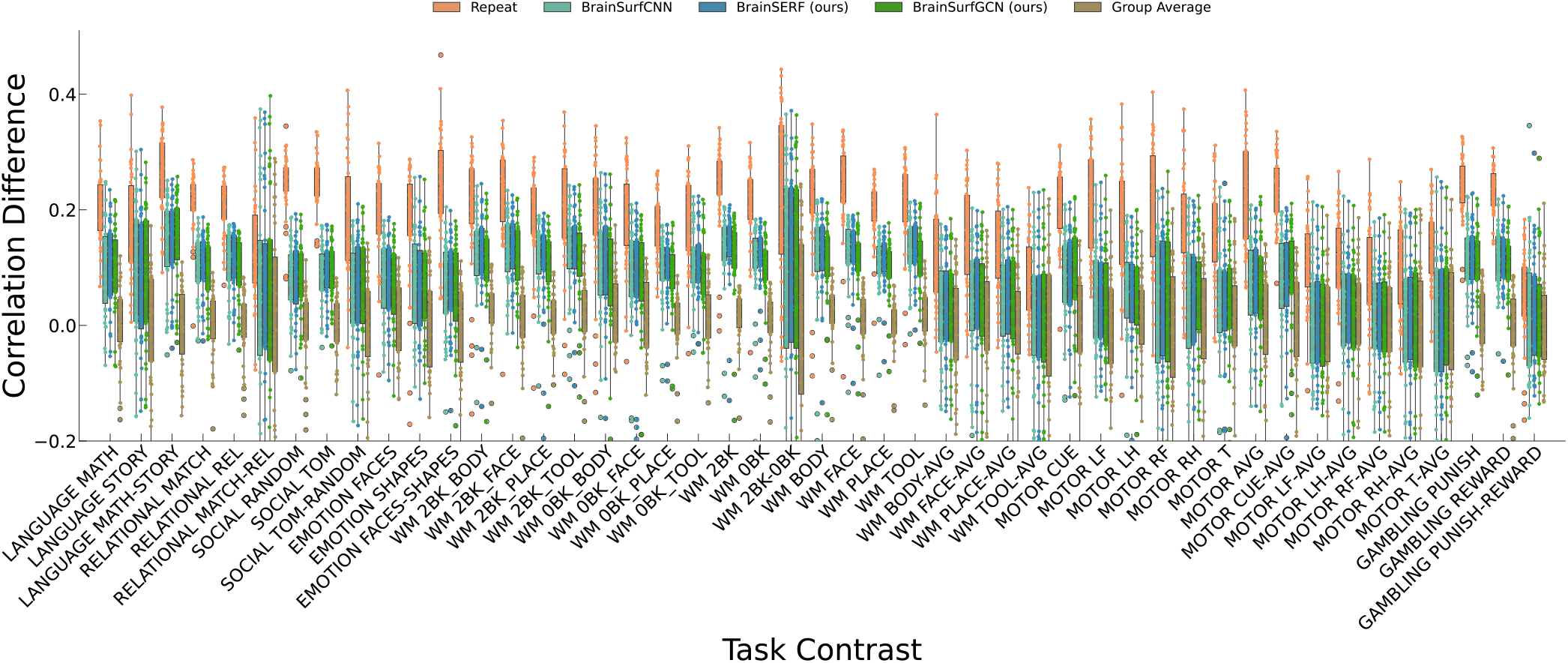
Difference in spatial correlation of predicted contrast to ground truth contrast of the same subject versus the mean spatial correlation of predicted contrast to ground truth contrasts of all other test set subjects. The correlation metric primarily measures the linear relationship between the model’s predicted task contrasts and the empirically observed (ground truth) contrasts. In this analysis, we derive the metric displayed in the figure by first calculating the correlation between a prediction based on a single subject’s resting-state data and the subject’s ground truth task activation contrast. Subsequently, we subtract the mean correlation of the same prediction from the task activation contrasts of all other subjects. Model predictions are compared against both the repeat condition and the group average baseline. A high correlation difference value signifies that the spatial correlation of a given prediction is substantially higher when comparing the prediction to the ground truth contrast of the same subject, as opposed to the ground truth contrasts of different subjects. This indicates that the model proficiently captures the idiosyncratic features of each individual’s task-related brain activity.

Next, we performed a subject identification task: for each predicted contrast, we computed its correlation with all ground-truth contrasts across the test set and evaluated whether the highest correlation matched the true subject. This classification-based metric offers a concrete measure of the identifiability of individuals from predicted brain maps. To examine the effect of input dimensionality, we systematically varied the number of ICA components (15, 25, 50, and 100 ICs). Average subject identification accuracies across task contrasts are summarized in Table 2.

**Table 2:** Subject Identification Test Accuracy Across Varying ICs.

| Model | 15 ICs | 25 ICs | 50 ICs | 100 ICs | Avg |
| --- | --- | --- | --- | --- | --- |
| BrainSurfCNN | 80.3 | 81.0 | 79.5 | 79.3 | 81.8 |
| BrainSERF (ours) | **80.6** | **82.3** | 80.4 | 79.8 | **82.8** |
| BrainSurfGCN (ours) | 79.5 | 79.5 | **80.7** | **79.9** | 82.4 |
| repeat (Benchmark) | **92.9** |  |  |  |  |

Interestingly, all models achieved comparable accuracy levels, but remained generally below the repeat benchmark. The average subject identification accuracy for repeat contrasts was 92.9% across 47 task contrasts, highlighting the strong test-retest reliability of the data. Notably, in a few tasks (e.g., Language: Math and Language: Story), model predictions slightly exceeded identifiability from repeat scans (S5)•

The performance did not consistently improve with increasing ICA dimensionality. Instead, subject identification accuracy remained relatively stable across resolutions, with no clear benefit beyond moderate dimensionality. This suggests that increasing input dimensionality alone does not necessarily translate to improved individual-level prediction.

### 3.3 Mechanisms of Variability in Predictability

Having established that all three architectures can generate individualized task activation patterns, we next asked why prediction accuracy varies across individuals and contrasts. We focused on three potential sources of variability grounded in cognitive neuroscience: task engagement, resting-state data quality, and inter-subject variability in task-evoked activation.

#### 3.3.1 Task Engagement Influences Predictability

We first examined whether behavioral task engagement modulates the success of rest-to-task prediction. To this end, we focused on the held-out test set (*n* = 39) and stratified participants based on their task performance (e.g., accuracy or behavioral variability). If a participant performs poorly on a task, reflected in lower accuracy, or greater behavioral variability, the neural representations associated with that task may be weaker or less consistently expressed, reducing the model’s ability to reconstruct individual activation patterns. For the behavioral performance analysis, we restricted our focus to a subset of 14 task contrasts that included valid accuracy metrics, allowing us to stratify subjects into high (above median) and low (below median) performance groups.

Although several contrasts showed nominal significance prior to correction, most effects did not survive multiple-comparison correction using the Benjamini-Hochberg FDR procedure. At the model level (Figure 7a), nominally significant group differences were observed for the repeat data and BrainSurfGCN, with above-average performers showing higher correlation differences than below-average performers; however, these effects did not survive FDR correction, potentially reflecting limited statistical power.

**Figure 7:**
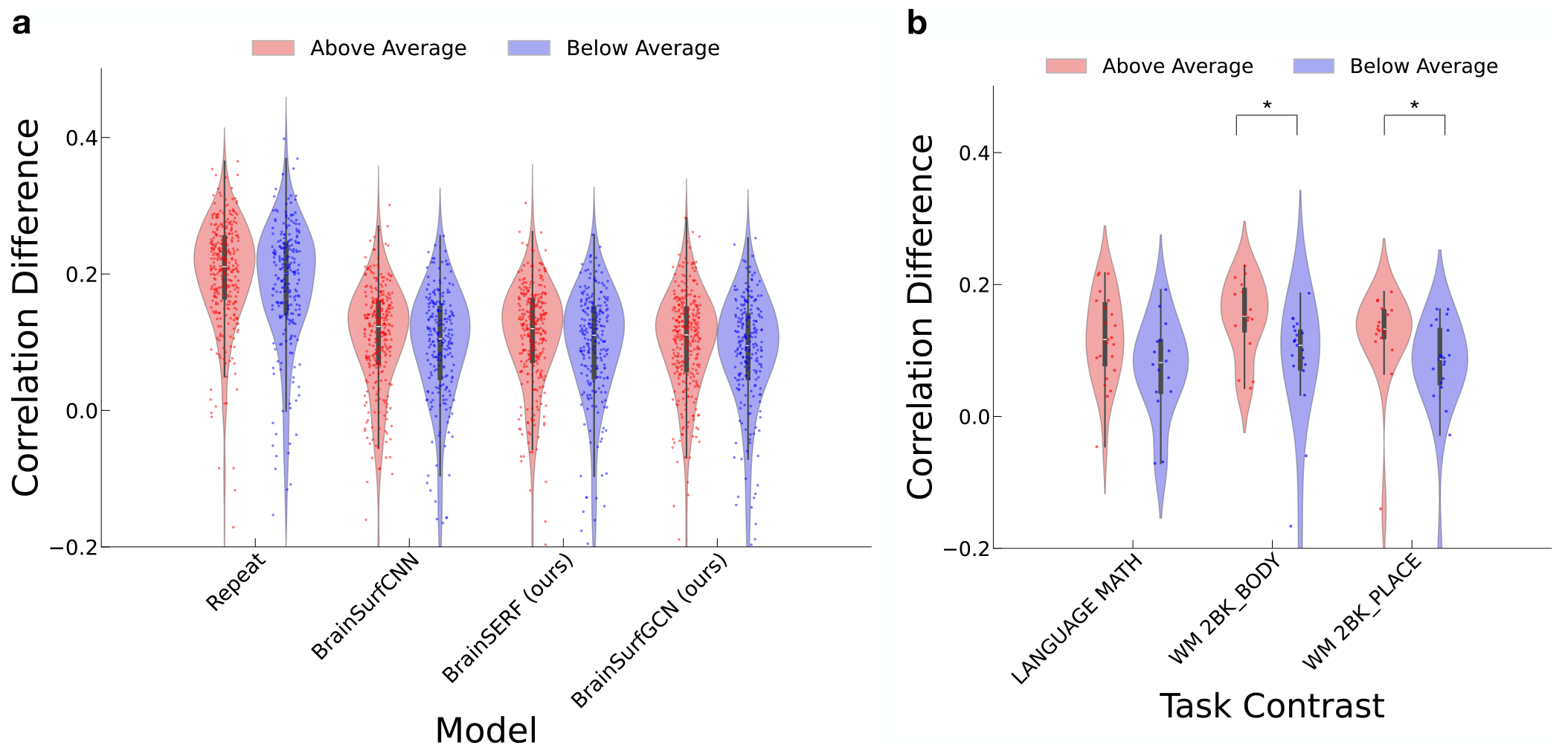
Correlation difference for above- and below-average task accuracy groups. Subjects were divided into above-average (red) and below-average (blue) behavioral performance groups for task contrasts with an available accuracy metric, and correlation differences were compared between groups. **(a)** Across models (repeat, BrainSurfCNN, BrainSERF, and BrainSurfGCN), no group differences remained significant after multiple-comparison correction (Benjamini-Hochberg FDR). Each point represents a subject’s correlation difference for a single task contrast (aggregated across all 14 task contrasts and 39 subjects; *n* = 546 points per model). **(b)** Three representative task contrasts are shown (BrainSurfGCN). Among all contrasts, only Working Memory: 2-Back Body and Working Memory: 2-Back Place showed significant group differences after FDR correction, with higher correlation differences in the above-average group. Here, each point represents a single subject (n = 39 per contrast).

Motivated by these nominal effects, we further examined task-specific patterns using BrainSurfGCN (Figure 7b). At the task level, only two contrasts, Working Memory: 2-Back Body and Working Memory: 2-Back Place, showed borderline significance after correction, with higher correlation differences in the above-average group.

All task contrasts are shown in Supplementary Figure S16, illustrating the full distribution of effects across tasks. Overall, these results suggest that the relationship between behavioral performance and predictability is limited, task-specific, and modest in magnitude. Accordingly, these analyses should be interpreted as exploratory and hypothesis-generating.

#### 3.3.2 Resting-State Signal Quality Shapes Prediction Error

We next examined whether resting-state data quality constrains prediction performance. Specifically, we evaluated the relationship between regional temporal signal-to-noise ratio (tSNR) and prediction error, measured as MSE.

We first computed the average tSNR across subjects for each cortical vertex (Figure 8a). Regions with lower tSNR, such as the orbitofrontal cortex and temporal poles, are known to be susceptible to susceptibility artifacts and physiological noise. We then compared the spatial distribution of low-tSNR regions with areas showing high prediction error, computed by comparing model predictions with repeat contrasts.

**Figure 8:**
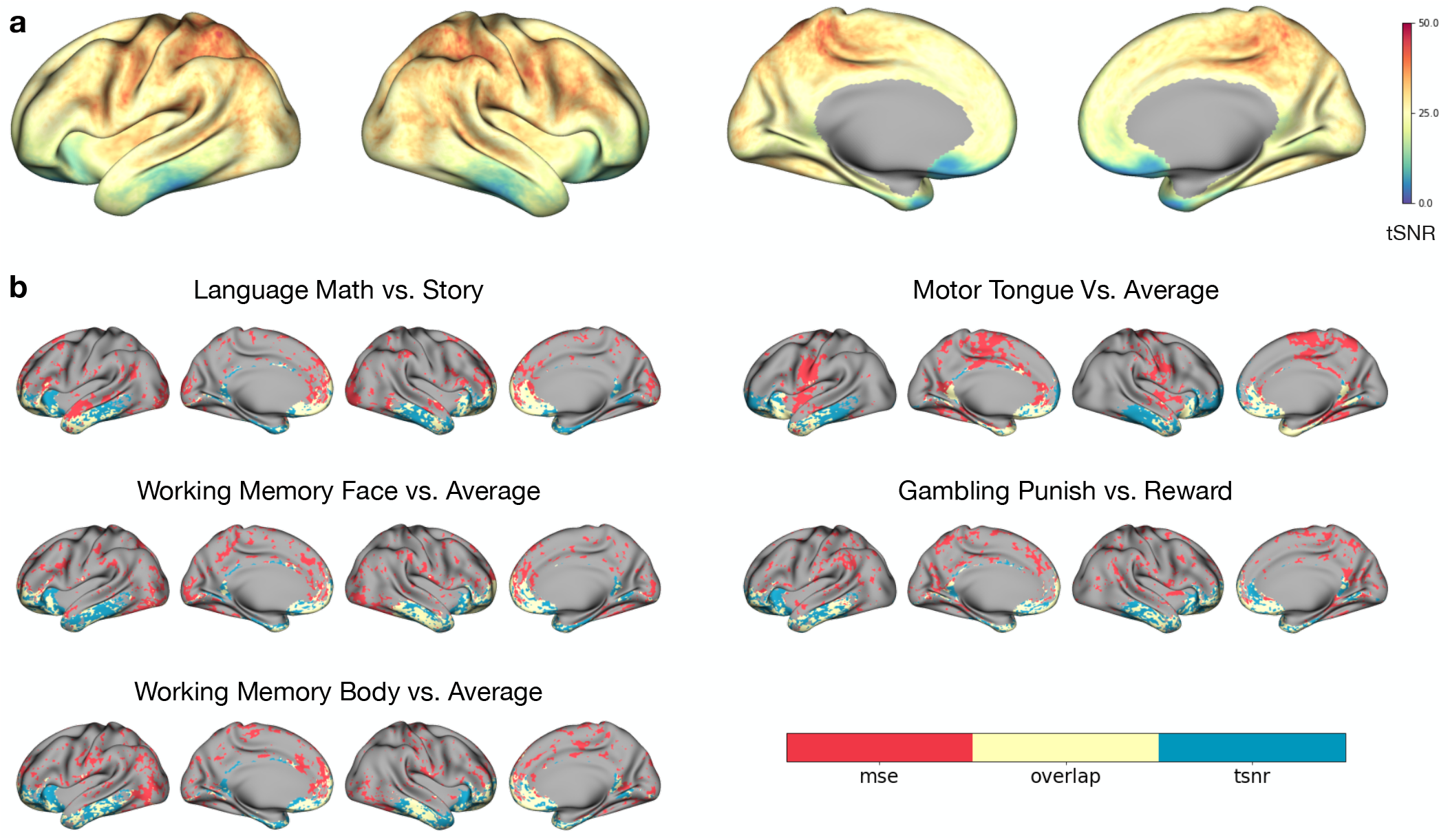
Relationship between lower tSNR and higher MSE of BrainSurfGCN’s predictions on the test set. **(a)** We computed the average tSNR for each vertex on the mesh across the test set and plotted it on the cortex. **(b)** The top 5 contrasts with the most overlap between the MSE and tSNR using the Dice AUC metric. We measured the overlap between the regions of the highest MSE and lowest tSNR. In this depiction, these maps are shown with a 25% threshold applied to both, allowing us to analyze the specific regions associated with the highest model error.

Across all 47 contrasts, delta contrasts (n=16) showed significantly higher error-to-tSNR Dice AUC than single-condition contrasts (n=31; mean 0.108 vs 0.088; one-sided Mann-Whitney U test, p=0.004; 10,000-permutation test, p=0.001; Cliff ‘s *δ* =0.48, large effect; Supplementary Figure S17). All five contrasts with the highest overlap were delta contrasts, including Language: Math - Story, Working Memory: Face - Avg, and Motor: Tongue - Avg. These results indicate that rest-to-task prediction performance is partly limited by the intrinsic noise characteristics of resting-state data, especially for contrasts requiring subtle condition-specific distinctions.

#### 3.3.3 Variability in Cortical Activation Across Subjects and Contrasts

Finally, we asked whether inter-subject variability in task-evoked activation constrains predictability. Prior studies have shown that contrasts involving subtle cognitive differences, particularly delta contrasts, exhibit less spatial consistency across individuals than single-condition contrasts (Seghier and Price, 2016).

To quantify this, we generated activation-frequency maps for each contrast by computing, for each subject, the top 10% of activated vertices and then calculating the proportion of subjects showing activation at each vertex. These maps were computed for the empirical data (test and repeat) and for predictions from all three models.

As illustrated in Figure 9, delta contrasts showed markedly greater spatial variability than single-condition contrasts. Even the repeat contrasts showed low spatial correlation for delta contrasts (e.g., Relational Match - Relational: r = 0.626), underscoring their low test-retest reliability. In contrast, single-condition contrasts such as Relational: Relational and Relational: Match showed high repeatability (r ≈ 0.97), reflecting stable and stereotyped activation patterns.

**Figure 9:**
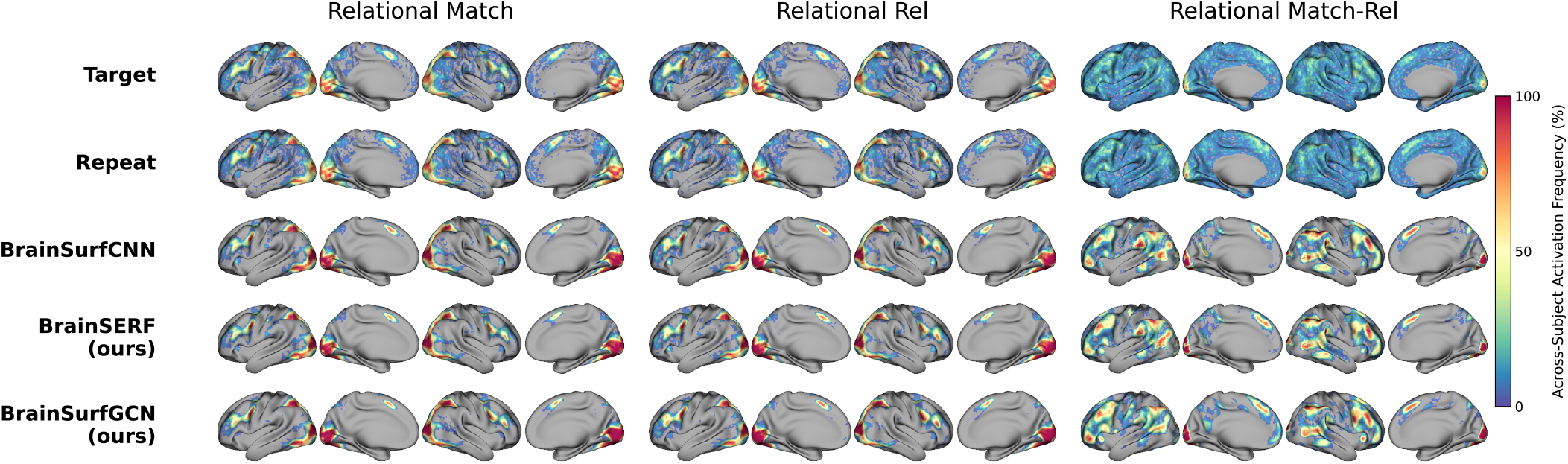
Cortical frequency plots associated with relational task contrasts. For each task contrast, we threshold at 10% and count the number of subjects that show activation on the vertex. This gives us a vertex value between 0 and 100, indicating the percentage of subjects with this vertex in the top 10% of activated vertices. Delta contrast maps indicating variability across the population show substantially greater spatial variation compared to both the repeat and the model-predicted contrasts.

The models reproduced these broad patterns: predictions were smoother and more spatially constrained than the empirical data and showed reduced correspondence for delta contrasts relative to single-condition contrasts. This suggests that high inter-subject variability, combined with lower reliability of delta contrasts, limits the upper bound of predictability for these contrasts, even for the empirical repeat maps.

Supplementary Tables S3 and S4 provide full spatial correlation matrices, further highlighting the dominant influence of the repeat-derived noise ceiling on contrast-level predictability.

Together, these analyses reveal that variability in rest-to-task prediction arises from a combination of cognitive factors (task engagement), technical factors (resting-state data quality), and neurobiological factors (inter-subject variability in cortical recruitment). These findings emphasize that prediction performance is not solely determined by model architecture, but is fundamentally shaped by the stability and reliability of the underlying neural signals.

## 4 Discussion

In this work, we reproduced the predictive performance of BrainSurfCNN and introduced two novel variants, including BrainSERF, which integrates a channel-wise attention mechanism for adaptive feature refinement, and BrainSurfGCN, which implements a graph-based representation to explicitly model the organization of brain networks. These models were designed in response to two central questions raised in the introduction: (1) Can deep learning models of resting-state to task prediction be made more architecturally efficient and fidelity? And (2) What are the key drivers of variability in individual-level prediction performance?

### 4.1 Model Innovations: Attention and Graph-based

#### Architectures

BrainSERF incorporates a channel-wise attention mechanism (SE block) that allows the model to adaptively scale the contribution of each IC map. This mimics a form of neural gain control, potentially introducing structured feature reweighting consistent with known neural gain modulation by enabling the model to emphasize or suppress distributed sources, analogous to dynamic reweighting in cortical circuits. While performance gains were modest, BrainSERF consistently achieved higher subject identification accuracy compared to the baseline.

BrainSurfGCN focused on computational efficiency, achieving more than 15 times reduction in trainable parameters without sacrificing performance. BrainSurfCNN and BrainSERF apply spherical convolution using parameterized differential operators that rely on four mesh representations: the original mesh, gradients along the *x* and *y* surface directions, and the Laplacian. While effective, this approach increases memory and compute demands by requiring multiple large tensors to be stored and processed at each layer. In contrast, our new BrainSurfGCN model uses a graph-based message-passing framework that represents the cortical surface as a node-edge graph and updates vertex features via a shared learnable weight matrix Θ. This design eliminates the need to explicitly compute or store differential operators, while still capturing local one-hop information propagation like spherical convolutions. The result is a dramatic reduction in model size (from ∼100MB to <1MB) and training time (from ∼26 to ∼6 hours), without sacrificing predictive performance. This efficiency makes BrainSurfGCN well-suited for resource-constrained environments, rapid prototyping, and potential clinical deployment, where model size and training speed are critical considerations. Although all models achieved comparable prediction accuracy, BrainSurfGCN substantially reduced computational cost, highlighting how graph structural designs may offer a scalable and efficient approach to individualized brain modeling.

### 4.2 Data Requirements and Model Scalability

A critical consideration in connectome fingerprinting approaches is the amount of data required to train predictive models. CNN-based models, including those evaluated in this study, typically involve a large number of trainable parameters and therefore benefit from large-scale datasets. In our case, over 900 subjects were used for training, which may limit the applicability of such models in smaller or more specialized datasets. In contrast, prior work using regression and penalized regression approaches has demonstrated strong predictive performance and subject-level specificity even with relatively modest sample sizes Osher et al., 2019; Tavor et al., 2016; Tobyne et al., 2018. These methods are therefore particularly attractive in data-limited settings.

Recent work has further highlighted alternative modeling strategies, including mechanistic models such as activity flow mapping Cole et al., 2016, as well as deep learning approaches that directly model spatial structure Ngo et al., 2022. These approaches highlight that prediction performance depends not only on model complexity but also on how connectivity information is represented and utilized. Within this broader landscape, BrainSurfGCN can be viewed as a middle ground between high-capacity deep learning models and more interpretable, low-parameter approaches. Notably, the reduction in model parameters in BrainSurfGCN is substantial (Table 1), suggesting a meaningful shift in the tradeoff between model capacity and scalability. By incorporating topology-aware structural constraints, it reduces model complexity while maintaining comparable predictive performance to CNN-based models. This suggests potential improvements in data efficiency and scalability, making it a promising alternative when large training datasets are not available. Taken together, these findings highlight that model choice should be guided not only by predictive performance but also by data availability, computational cost, and the degree of structural inductive bias incorporated into the model.

### 4.3 Positioning CNN-Based Models Within Connectome Fingerprinting Approaches

A growing body of work has demonstrated that task-evoked brain activity can be predicted from patterns of structural or functional connectivity using a range of modeling approaches (Cole et al., 2016; Ngo et al., 2022; Tavor et al., 2016). These approaches differ substantially in their underlying assumptions and modeling strategies. Regression-based methods provide interpretable mappings between connectivity features and task activation, and have been shown to capture strong subject-level specificity. Modeling approaches such as activity flow mapping Cole et al., 2016 provide an explicit account of how task-evoked activity may arise from the propagation of activity over intrinsic connectivity networks. In contrast, CNN-based models Ngo et al., 2022 are primarily optimized to reconstruct spatially structured activation patterns, often capturing spatial distributions effectively, but without explicitly modeling the underlying network topology. Prior work by Ngo et al., 2022 showed that CNN-based models achieve performance broadly comparable to both group-average baselines and linear regression approaches in terms of Dice AUC, with only modest differences across methods. However, these differences are relatively small, indicating that a substantial portion of task-evoked activation patterns can already be captured by simpler models.

Importantly, predictive performance across all approaches is fundamentally constrained by the reliability of the underlying data, as reflected by repeat measurements, which serve as a practical noise ceiling. Notably, the gap between model predictions and repeat data is substantially larger than the gap between different modeling approaches, underscoring the dominant role of intrinsic variability in limiting prediction accuracy. Within this broader landscape, our work focuses on how architectural design choices influence the trade-off between predictive performance, computational efficiency, and representational structure.

Specifically, BrainSERF introduces channel-wise modulation mechanisms that enable adaptive feature reweighting, while BrainSurfGCN explicitly incorporates cortical topology to constrain information flow.

Unlike conventional CNN-based approaches that rely on local convolutional operations without explicitly modeling networks, BrainSurfGCN incorporates connectivity-informed message passing to constrain information flow. This design enables the model to capture interactions between distributed brain regions while maintaining comparable predictive performance with reduced computational cost. These findings suggest that embedding structured priors into model architecture offers an effective strategy for improving efficiency and interpretability in rest-to-task prediction.

### 4.4 Impact of Input Representation and Dimensionality

Moreover, we examined the role of representational dimensionality by varying the number of ICA components used as model input. Contrary to prior findings that higher ICA dimensionality can improve prediction performance (Tavor et al., 2016; Tobyne et al., 2018), we observed that performance remained relatively stable across a wide range of component numbers. This suggests that predictive information may saturate at relatively low-dimensional representations in this setting.

One possible explanation is that the model architectures already capture relevant spatial structure through learned filters, reducing the marginal benefit of increasing input dimensionality. Alternatively, increasing ICA dimensionality may introduce redundant or noisy features that do not contribute additional predictive signal. It is also possible that prediction performance is fundamentally constrained by the reliability of task contrasts, such that improvements in input representation yield diminishing returns beyond a certain point. This may also indicate that rest-to-task prediction is limited more by the information content of the input data than by its dimensionality.

### 4.5 Understanding variability in model performance

Our second line of investigation focused on understanding why model performance varies across task contrasts and individuals. We observed that prediction accuracy was influenced by task engagement, data quality, and inter-subject variability.

First, we found preliminary evidence that poorer model predictions were associated with lower task accuracy and slower reaction times, particularly in tasks requiring sustained attention or complex reasoning. This relationship between poor prediction performance and poor task performance has also been shown by Gonzalez-Castillo et al., 2015, where the prediction of cognitive states from functional connectivity was impaired by a possible loss of concentration or awareness. We present preliminary evidence for our hypothesis; rigorous future work is needed to better account for task performance during prediction.

Notably, the observed relationships between prediction accuracy and behavioral performance were modest in magnitude. This is consistent with prior work showing that task-fMRI measures have limited test-retest reliability, and that behavioral performance reflects multiple cognitive processes that are only partially captured by large-scale activation patterns. Additionally, variability in attention and engagement during both resting-state and task scans may further attenuate these relationships. These findings should therefore be interpreted as preliminary indicators of sources of variability, rather than strong predictive links. This further suggests that variability in prediction performance may be driven more by underlying neural and behavioral variability than by model differences alone.

Second, we investigated why delta contrast maps (i.e., condition A vs. condition B) yielded lower prediction and test-retest reliability compared to contrasts versus baseline. An exploratory analysis of repeat error and tSNR maps revealed that delta contrasts showed greater overlap between signal dropout regions (low tSNR) and areas of high prediction error, particularly in contrasts involving subtle task manipulations (e.g., Math vs. Story, Punish vs. Reward). This suggests that delta contrasts may reflect noisier or less stable signal differences, making them more challenging targets for prediction models. Future studies should consider modeling task contrasts with varying levels of reliability separately or weighting voxels based on tSNR-informed confidence.

Third, we examined the role of inter-subject variability in shaping prediction performance. As illustrated in Figure 9, the degree of spatial consistency varied substantially across task contrasts. Regions associated with more stimulus-driven processing, such as visual areas, showed relatively high consistency across subjects, with activation patterns concentrated in similar locations. In contrast, less predictable contrasts, particularly delta contrasts, exhibited more diffuse and spatially variable activation patterns across individuals. This variability was also reflected in repeat measurements, indicating that it is not specific to model predictions. Together, these findings suggest that inter-subject variability in cortical activation, and the extent to which specific regions are consistently engaged across individuals, plays a key role in determining prediction performance.

### 4.6 Limitations and future direction

Our study has several limitations, spanning both methodological design and generalization, that open avenues for future research: (1) Architectural scope: While BrainSERF and BrainSurfGCN introduced efficient design changes, the performance gains were incremental. New architectural innovations, such as equivariant GNNs, spectral attention, or hybrid encoding-decoding frameworks (Kondor, 2025), may yield greater improvements in both accuracy and interpretability. (2) Generalization across mesh structures: BrainSurfGCN was evaluated on a fixed cortical mesh; however, its flexibility to operate on graphs of varying resolution remains untested. Leveraging coarser meshes, as done in the original BrainSurfCNN, may offer further speedups with minimal performance loss. (3) Zero-shot task generalization: Our models were trained to predict 47 fixed task contrasts. Extending these models to unseen tasks remains an open challenge and may require task embedding approaches or transfer learning on task instructions. (4) Group IC derivation: Resting-state features were constructed using group-level IC maps provided by HCP, which may include some overlap with test subjects. Although this risk is minimal given the dataset size, future work should evaluate whether using subject-specific or held-out group ICs changes prediction fidelity. (5) Reliability of delta contrasts: Lower prediction performance for delta contrasts likely reflects their reduced reliability and higher noise sensitivity, a limitation consistently observed across prior work Tavor et al., 2016; Tripathi and Somers, 2023. This suggests that performance limits are driven more by intrinsic variability than by model capacity. (6) Cross-dataset generalization: All models were evaluated on HCP, limiting assessment of generalizability across datasets with different acquisition and population characteristics Ngo et al., 2022; Tik et al., 2023. (7) Family structure: The HCP dataset includes related individuals, and although standard splits were used, potential dependencies between training and test sets may slightly inflate performance estimates.

### 4.7 Toward efficient and scalable predictive modeling

Together, our findings highlight the importance of structurally informed modeling choices that enable efficient and scalable prediction of task-evoked brain activity from resting-state connectivity. In particular, BrainSurfGCN demonstrates that comparable predictive performance can be achieved with substantially reduced model complexity, emphasizing the value of parameter-efficient architectures for large-scale neuroimaging applications.

These results further support the idea that resting-state connectivity contains latent signatures of task-evoked activity, and that appropriately designed models can extract these signals in a subject-specific manner.

Moving forward, linking these predictive representations to behavior, trait variability, and clinical outcomes will be essential for translating these approaches into practical neuroscience and clinical settings. In addition, improving data quality, contrast reliability, and representation design will be critical for enhancing the robustness and generalizability of future models.

## Data and Code Availability

https://github.com/braindynamicslab All code used for this article is made publicly available on GitHub at https://github.com/braindynamicslab/dl-task-contrast-prediction

All data used in this article is publicly available through the Human Connectome Project.

## Acknowledgments

This work was supported by an NIH R01 MH127608 and an MCHRI Faculty Scholar Award to Manish Saggar. Data were provided by the Human Connectome Project, WU-Minn Consortium (Principal Investigators: David Van Essen and Kamil Ugurbil; 1U54MH091657) funded by the 16 NIH Institutes and Centers that support the NIH Blueprint for Neuroscience Research; and by the McDonnell Center for Systems Neuroscience at Washington University.

## Author Contributions

Soren Madsen: Conceptualization, methodology, software, validation, formal analysis, investigation, data curation, writing of original draft, reviewing and editing, and visualization. Young-Eun Lee: Methodology, software, validation, reviewing and editing, and visualization. Shaun K. L. Quah: Reviewing and editing. Lucina Q. Uddin: Conceptualization, reviewing and editing, and funding acquisition. Jeanette A. Mumford: Conceptualization, reviewing and editing, and funding acquisition. Deanna M. Barch: Conceptualization, reviewing and editing, and funding acquisition. Damien A. Fair: Conceptualization, reviewing and editing, and funding acquisition. Ian H. Gotlib: Conceptualization, reviewing and editing, and funding acquisition. Russell A. Poldrack: Conceptualization, reviewing and editing, supervision, and funding acquisition. Amy Kuceyeski: Conceptualization, validation, reviewing and editing, supervision, and funding acquisition. Manish Saggar: Conceptualization, methodology, investigation, resources, reviewing and editing, supervision, and funding acquisition.

## Competing Interests

No competing interests are present among the authors of this work.

## Footnotes

1 GitHub Link: https://github.com/ngohgia/brain-surf-cnn

## Notes

### Competing Interest Statement

The authors have declared no competing interest.

### Summary of Updates

We have carefully revised the manuscript. In particular, we have strengthened the framing of the work to more explicitly emphasize its neuroscientific contributions, clarified the positioning of our models relative to prior approaches, and expanded the analyses and discussion to better contextualize the results.

