## Supplementary Information for "Efficient Deep Learning Models for Predicting Individualized Task Activation from Resting-State Functional Connectivity"

### A Supplementary Information

#### A.1 Additional Results

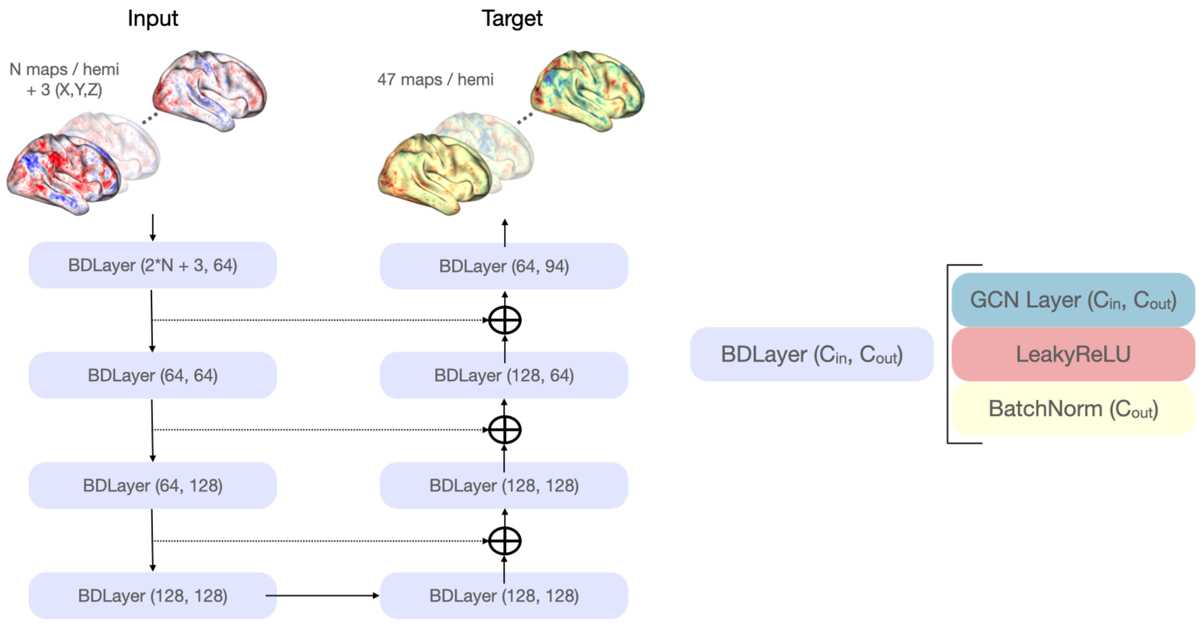


Figure S1: **A high-level overview of the BrainSurfGCN model architecture.** We chose this structure for the BDLayers of the model since the literature has shown that the combination of Graph Convolution, LeakyReLU, and BatchNorm performs well. There are 8 modules of BDLayers to ensure that information is disseminated across the entire mesh during one pass of the model.

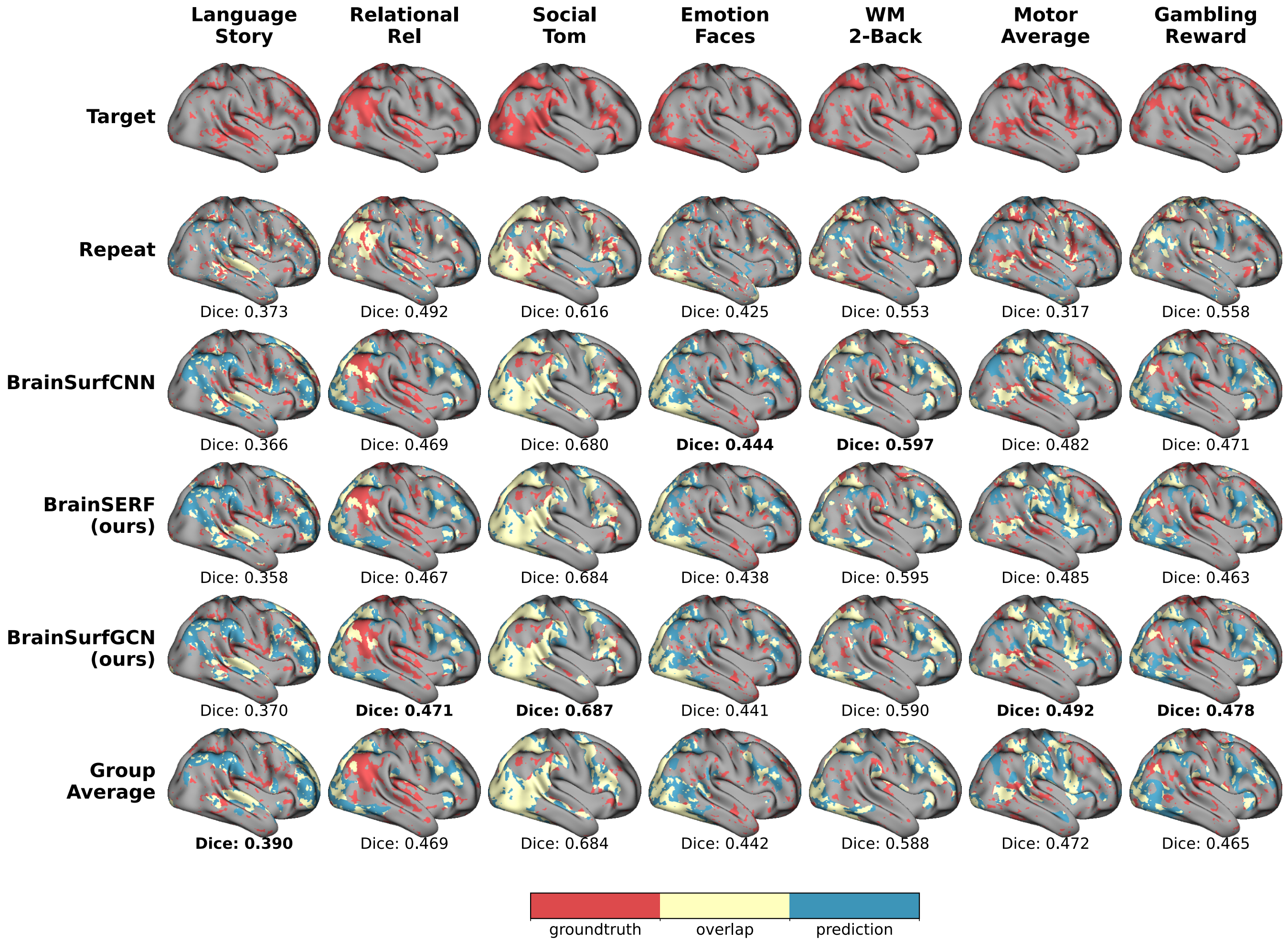


Figure S2: **Thresholded task activation maps for 7 task contrasts using threshold of 25% and IC of 15.** We compare ground truth, repeat, group average, and models' predicted activation for an individual (subject 917255). The thresholded activation maps on the right hemisphere (lateral view) represent, respectively, Language: Story, Relational Processing: Relational, Social Cognition: Theory of Mind, Emotion: Faces, Working Memory: 2-Back, Motor: Average, and Gambling: Reward. Ground truth (target), repeat scans, group average, and predictions from BrainSurfCNN, BrainSERF, and BrainSurfGCN are presented with colors indicating ground truth, model prediction, and overlap. Dice scores are displayed beneath each row, quantifying prediction fidelity across thresholds. The highest Dice value within each column is bolded to highlight the best-performing models.

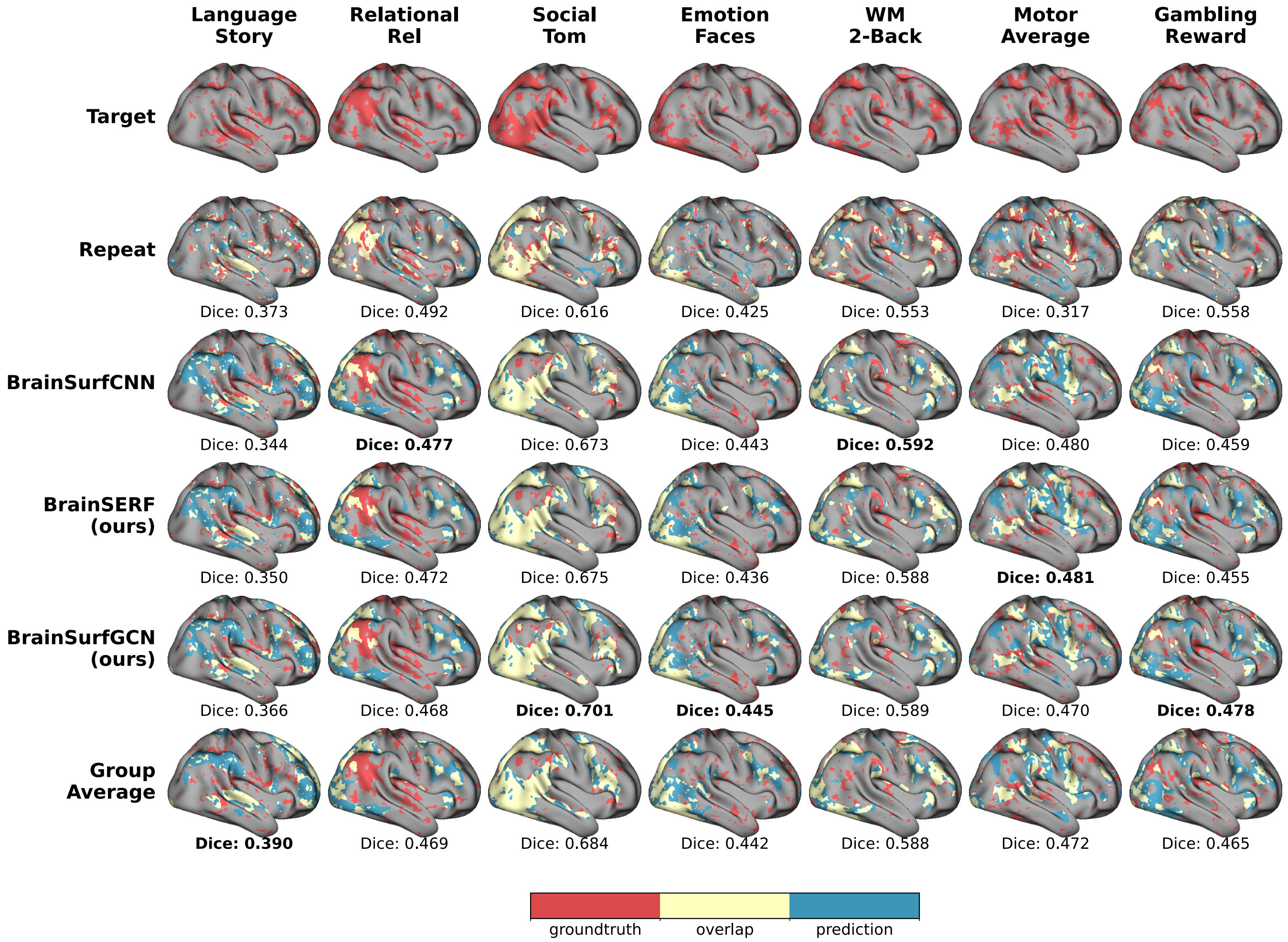


Figure S3: **Thresholded task activation maps for 7 task contrasts using threshold of 25% and IC of 25.** We compare ground truth, repeat, group average, and models' predicted activation for an individual (subject 917255). The thresholded activation maps on the right hemisphere (lateral view) represent, respectively, Language: Story, Relational Processing: Relational, Social Cognition: Theory of Mind, Emotion: Faces, Working Memory: 2-Back, Motor: Average, and Gambling: Reward. Ground truth (target), repeat scans, group average, and predictions from BrainSurfCNN, BrainSERF, and BrainSurfGCN are presented with colors indicating ground truth, model prediction, and overlap. Dice scores are displayed beneath each row, quantifying prediction fidelity across thresholds. The highest Dice value within each column is bolded to highlight the best-performing models.

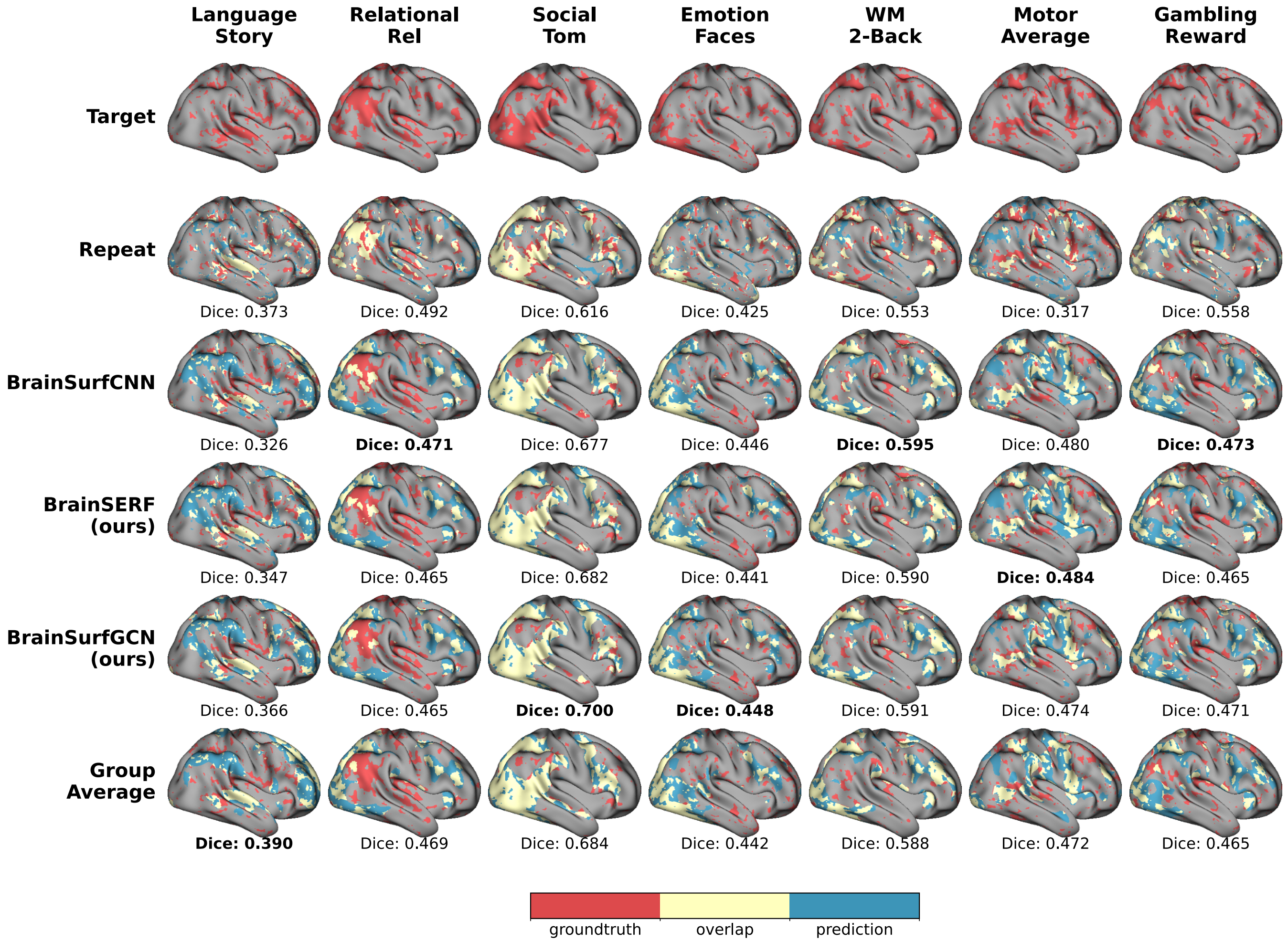


Figure S4: **Thresholded task activation maps for 7 task contrasts using threshold of 25% and IC of 100.** We compare ground truth, repeat, group average, and models' predicted activation for an individual (subject 917255). The thresholded activation maps on the right hemisphere (lateral view) represent, respectively, Language: Story, Relational Processing: Relational, Social Cognition: Theory of Mind, Emotion: Faces, Working Memory: 2-Back, Motor: Average, and Gambling: Reward. Ground truth (target), repeat scans, group average, and predictions from BrainSurfCNN, BrainSERF, and BrainSurfGCN are presented with colors indicating ground truth, model prediction, and overlap. Dice scores are displayed beneath each row, quantifying prediction fidelity across thresholds. The highest Dice value within each column is bolded to highlight the best-performing models.

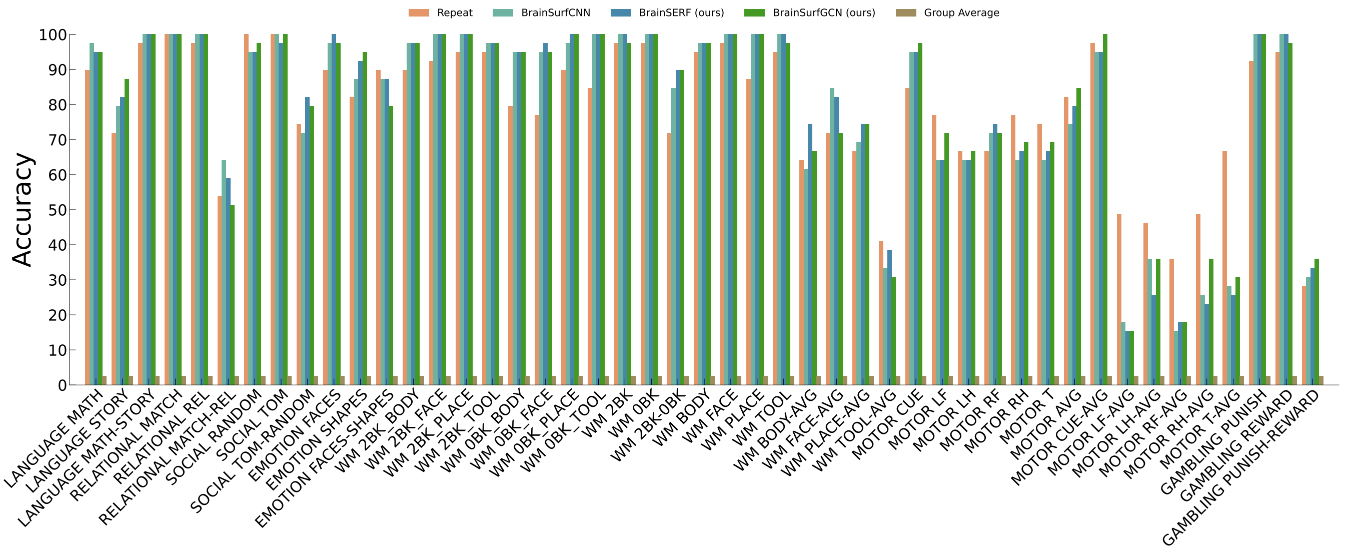


Figure S5: Subject identification accuracy across all task contrasts.

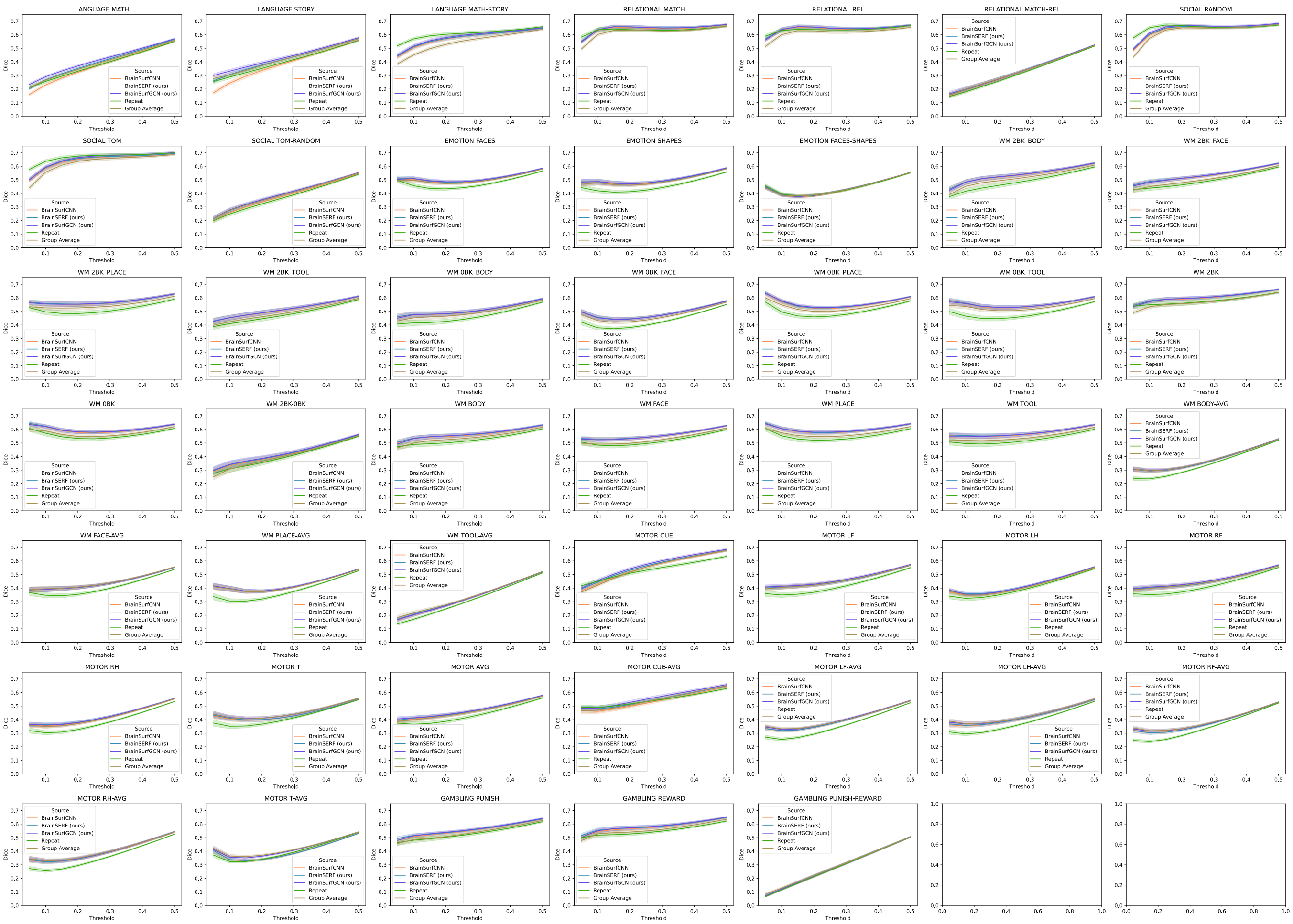


Figure S6: Dice scores of all models across all task contrasts.

Table S1: Dice AUC Across All Language, Relational, Social, Emotional, and Working Memory Tasks

| Contrast | Repeat | BrainSurfCNN | BrainSERF | BrainSurfGCN | Group Average |
|---|---|---|---|---|---|
| LANGUAGE MATH | 0.173 | 0.169 | 0.181 | **0.187** | 0.178 |
| LANGUAGE STORY | 0.181 | 0.174 | 0.190 | **0.196** | 0.189 |
| LANGUAGE MATH-STORY | **0.274** | 0.261 | 0.262 | 0.264 | 0.246 |
| RELATIONAL MATCH | 0.286 | 0.290 | **0.292** | **0.292** | 0.281 |
| RELATIONAL REL | 0.287 | 0.289 | **0.291** | **0.291** | 0.279 |
| RELATIONAL MATCH-REL | 0.145 | 0.150 | **0.151** | **0.151** | **0.151** |
| SOCIAL RANDOM | **0.295** | 0.289 | 0.290 | 0.293 | 0.284 |
| SOCIAL TOM | **0.302** | 0.292 | 0.293 | 0.296 | 0.284 |
| SOCIAL TOM-RANDOM | 0.168 | 0.173 | 0.178 | **0.180** | 0.178 |
| EMOTION FACES | 0.214 | 0.229 | 0.230 | **0.231** | 0.227 |
| EMOTION SHAPES | 0.205 | 0.225 | 0.226 | **0.227** | 0.223 |
| EMOTION FACES-SHAPES | 0.197 | 0.196 | **0.198** | 0.197 | **0.198** |
| WM 2BK_BODY | 0.220 | 0.242 | **0.243** | **0.243** | 0.232 |
| WM 2BK_FACE | 0.223 | 0.241 | **0.242** | **0.242** | 0.229 |
| WM 2BK_PLACE | 0.232 | 0.257 | **0.258** | **0.258** | 0.246 |
| WM 2BK_TOOL | 0.216 | 0.232 | **0.233** | **0.233** | 0.222 |
| WM 0BK_BODY | 0.209 | 0.229 | **0.230** | **0.230** | 0.222 |
| WM 0BK_FACE | 0.194 | 0.218 | **0.219** | 0.218 | 0.210 |
| WM 0BK_PLACE | 0.225 | 0.249 | 0.251 | **0.252** | 0.240 |
| WM 0BK_TOOL | 0.218 | 0.248 | **0.249** | **0.249** | 0.239 |
| WM 2BK | 0.261 | 0.272 | **0.273** | **0.273** | 0.258 |
| WM 0BK | 0.251 | 0.269 | **0.271** | **0.271** | 0.258 |
| WM 2BK-0BK | 0.182 | 0.189 | 0.191 | **0.192** | 0.184 |
| WM BODY | 0.237 | 0.254 | **0.255** | **0.255** | 0.244 |
| WM FACE | 0.233 | 0.250 | **0.251** | **0.251** | 0.238 |
| WM PLACE | 0.247 | 0.269 | 0.269 | **0.270** | 0.257 |
| WM TOOL | 0.236 | 0.257 | **0.259** | 0.258 | 0.245 |
| WM BODY-AVG | 0.156 | 0.168 | 0.169 | **0.171** | 0.170 |
| WM FACE-AVG | 0.183 | 0.199 | 0.199 | 0.199 | **0.200** |
| WM PLACE-AVG | 0.171 | 0.191 | 0.191 | **0.192** | 0.190 |
| WM TOOL-AVG | 0.141 | 0.150 | 0.151 | 0.150 | **0.153** |

Table S2: Dice AUC Across All Motor and Gambling Tasks

| Contrast | Repeat | BrainSurfCNN | BrainSERF | BrainSurfGCN | Group Average |
|---|---|---|---|---|---|
| MOTOR CUE | 0.241 | 0.250 | 0.254 | **0.258** | 0.249 |
| MOTOR LF | 0.189 | 0.207 | 0.207 | **0.210** | 0.207 |
| MOTOR LH | 0.181 | 0.188 | **0.191** | **0.191** | 0.188 |
| MOTOR RF | 0.189 | 0.204 | 0.204 | **0.208** | 0.204 |
| MOTOR RH | 0.173 | 0.190 | **0.192** | **0.192** | 0.189 |
| MOTOR T | 0.188 | 0.199 | 0.199 | **0.204** | 0.203 |
| MOTOR AVG | 0.196 | 0.209 | 0.211 | **0.213** | 0.209 |
| MOTOR CUE-AVG | 0.245 | 0.242 | 0.247 | **0.253** | 0.246 |
| MOTOR LF-AVG | 0.161 | 0.179 | 0.179 | **0.182** | **0.182** |
| MOTOR LH-AVG | 0.173 | 0.190 | 0.191 | 0.193 | **0.194** |
| MOTOR RF-AVG | 0.157 | 0.172 | 0.171 | 0.174 | **0.175** |
| MOTOR RH-AVG | 0.161 | 0.179 | 0.180 | **0.181** | **0.181** |
| MOTOR T-AVG | 0.179 | 0.178 | 0.178 | 0.187 | **0.189** |
| GAMBLING PUNISH | 0.239 | 0.250 | 0.252 | **0.253** | 0.242 |
| GAMBLING REWARD | 0.248 | 0.261 | 0.263 | **0.264** | 0.253 |
| GAMBLING PUNISH-REWARD | 0.128 | 0.129 | 0.130 | 0.130 | **0.131** |

Table S3: Spatial correlation of percentage of subjects showing top 10% activation between predicted and ground truth contrasts associated language, relational, social, emotional, and working memory tasks.

| Contrast | Repeat | BrainSurfCNN | BrainSERF | BrainSurfGCN | Group Average |
|---|---|---|---|---|---|
| LANGUAGE MATH | 0.256 | 0.231 | 0.268 | **0.291** | 0.265 |
| LANGUAGE STORY | 0.289 | 0.244 | 0.307 | **0.332** | 0.304 |
| LANGUAGE MATH-STORY | **0.570** | 0.505 | 0.512 | 0.516 | 0.452 |
| RELATIONAL MATCH | **0.637** | 0.633 | **0.637** | **0.637** | 0.602 |
| RELATIONAL REL | 0.632 | 0.636 | **0.640** | 0.639 | 0.601 |
| RELATIONAL MATCH-REL | 0.180 | 0.192 | 0.195 | **0.199** | **0.199** |
| SOCIAL RANDOM | **0.652** | 0.598 | 0.605 | 0.614 | 0.575 |
| SOCIAL TOM | **0.637** | 0.585 | 0.591 | 0.594 | 0.557 |
| SOCIAL TOM-RANDOM | 0.249 | 0.259 | 0.274 | **0.279** | 0.276 |
| EMOTION FACES | 0.457 | 0.506 | 0.509 | **0.510** | 0.498 |
| EMOTION SHAPES | 0.419 | 0.482 | 0.486 | **0.488** | 0.479 |
| EMOTION FACES-SHAPES | **0.395** | 0.388 | 0.393 | 0.388 | 0.388 |
| WM 2BK_BODY | 0.416 | **0.485** | 0.484 | 0.483 | 0.453 |
| WM 2BK_FACE | 0.440 | 0.486 | **0.489** | 0.482 | 0.451 |
| WM 2BK_PLACE | 0.497 | 0.558 | **0.559** | 0.555 | 0.526 |
| WM 2BK_TOOL | 0.410 | 0.453 | **0.454** | 0.453 | 0.423 |
| WM 0BK_BODY | 0.416 | 0.477 | **0.480** | 0.477 | 0.453 |
| WM 0BK_FACE | 0.379 | 0.454 | **0.456** | 0.453 | 0.433 |
| WM 0BK_PLACE | 0.496 | 0.570 | 0.574 | **0.578** | 0.548 |
| WM 0BK_TOOL | 0.466 | 0.557 | **0.562** | 0.558 | 0.537 |
| WM 2BK | 0.550 | 0.576 | **0.578** | 0.573 | 0.534 |
| WM 0BK | 0.572 | 0.619 | **0.623** | 0.621 | 0.589 |
| WM 2BK-0BK | 0.309 | 0.334 | 0.340 | **0.342** | 0.307 |
| WM BODY | 0.490 | 0.534 | **0.536** | 0.533 | 0.500 |
| WM FACE | 0.484 | 0.527 | **0.529** | 0.523 | 0.492 |
| WM PLACE | 0.553 | 0.604 | **0.607** | 0.606 | 0.576 |
| WM TOOL | 0.496 | 0.551 | **0.555** | 0.550 | 0.520 |
| WM BODY-AVG | 0.236 | 0.293 | 0.293 | **0.299** | 0.295 |
| WM FACE-AVG | 0.347 | 0.390 | 0.392 | 0.392 | **0.395** |
| WM PLACE-AVG | 0.305 | 0.394 | **0.396** | **0.396** | 0.394 |
| WM TOOL-AVG | 0.170 | 0.201 | 0.207 | 0.203 | **0.217** |

Table S4: Spatial correlation of percentage of subjects showing top 10% activation between predicted and ground truth contrast sets associated with motor and gambling tasks.

| Contrast | Repeat | BrainSurfCNN | BrainSERF | BrainSurfGCN | Group Average |
|---|---|---|---|---|---|
| MOTOR CUE | 0.450 | 0.430 | 0.444 | **0.455** | 0.422 |
| MOTOR LF | 0.349 | 0.407 | 0.406 | **0.414** | 0.405 |
| MOTOR LH | 0.324 | 0.348 | **0.358** | 0.354 | 0.349 |
| MOTOR RF | 0.352 | 0.400 | 0.402 | **0.408** | 0.398 |
| MOTOR RH | 0.303 | 0.356 | **0.363** | 0.360 | 0.349 |
| MOTOR T | 0.352 | 0.405 | 0.410 | **0.416** | 0.414 |
| MOTOR AVG | 0.366 | 0.405 | 0.411 | **0.414** | 0.399 |
| MOTOR CUE-AVG | **0.488** | 0.461 | 0.479 | 0.484 | 0.470 |
| MOTOR LF-AVG | 0.254 | 0.317 | 0.319 | 0.328 | **0.332** |
| MOTOR LH-AVG | 0.295 | 0.355 | 0.358 | 0.369 | **0.371** |
| MOTOR RF-AVG | 0.238 | 0.305 | 0.303 | 0.315 | **0.318** |
| MOTOR RH-AVG | 0.255 | 0.323 | 0.322 | 0.328 | **0.329** |
| MOTOR T-AVG | 0.324 | 0.337 | 0.338 | 0.356 | **0.364** |
| GAMBLING PUNISH | 0.481 | 0.508 | **0.516** | 0.513 | 0.487 |
| GAMBLING REWARD | 0.518 | 0.545 | **0.555** | 0.550 | 0.528 |
| GAMBLING PUNISH-REWARD | 0.116 | 0.119 | 0.120 | 0.121 | **0.128** |

#### A.2 Examples of Unthresholded Task Contrast Prediction

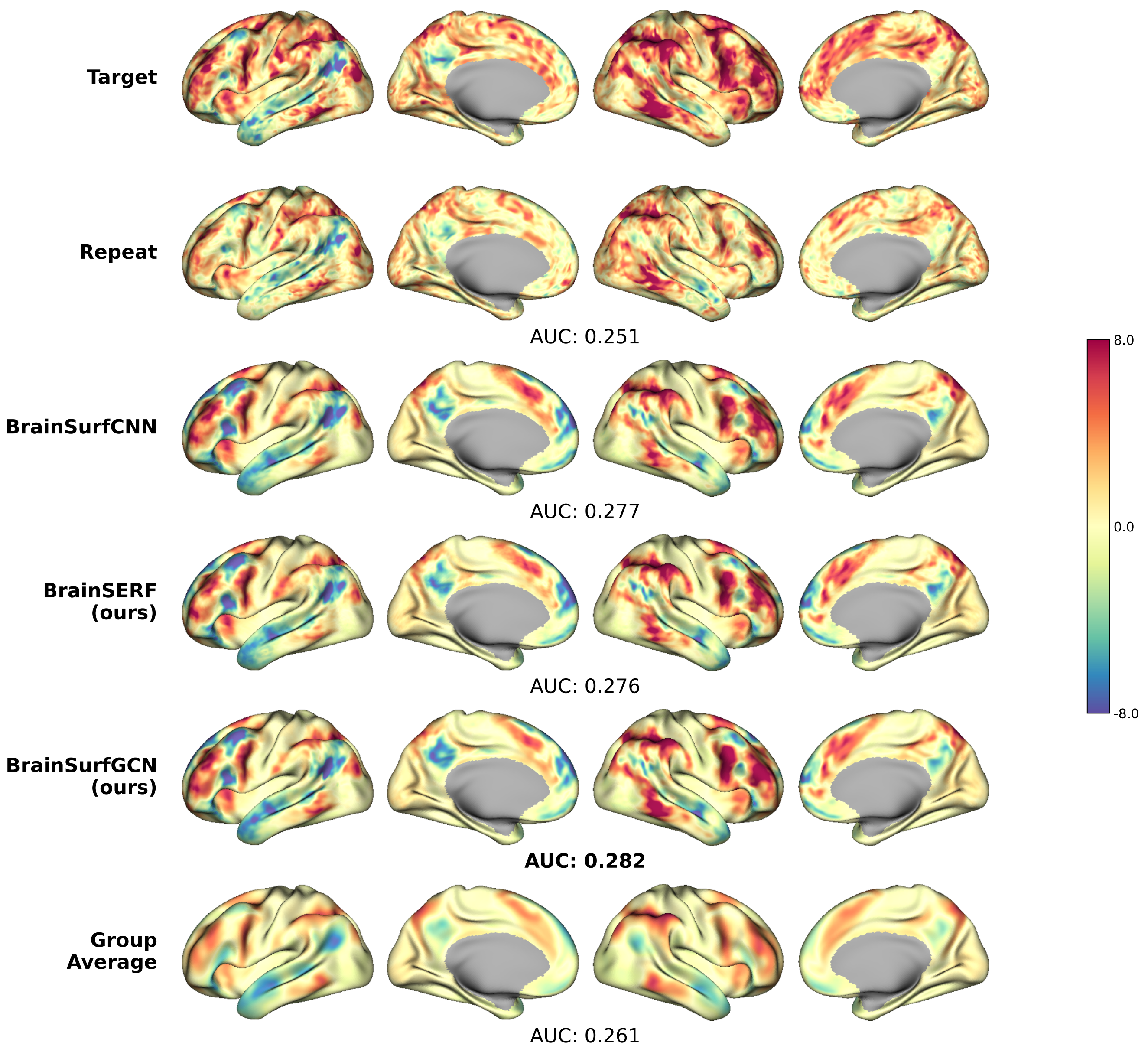


Figure S7: **Examples of unthresholded task activation for Language: Math-Story.** We compare ground truth (target), repeat, model-predicted activation, and group average for an individual (subject 917255).

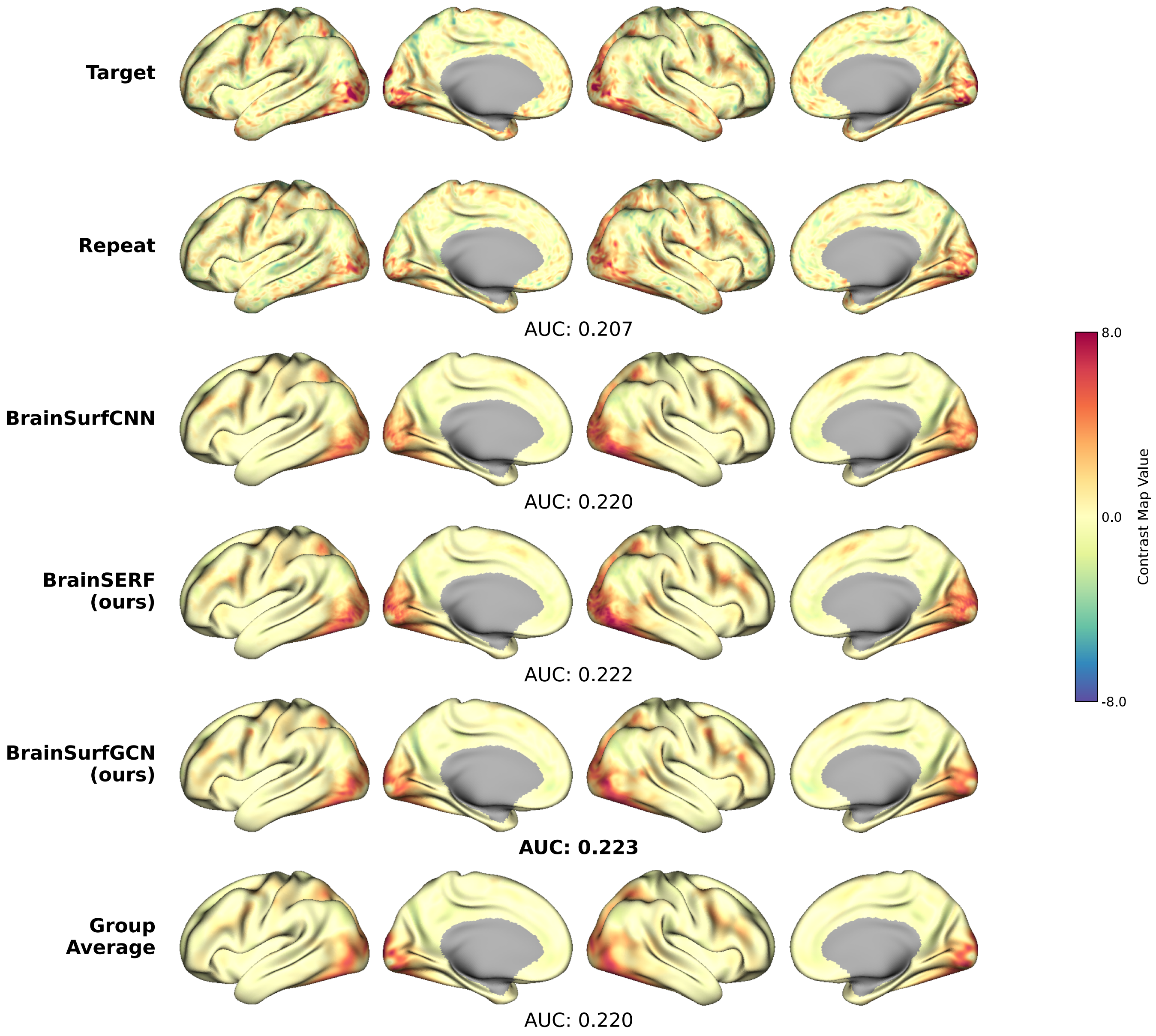


Figure S8: **Examples of unthresholded task activation for Emotion: Face.** We compare ground truth (target), repeat, model-predicted, and group average activation for an individual (subject 917255).

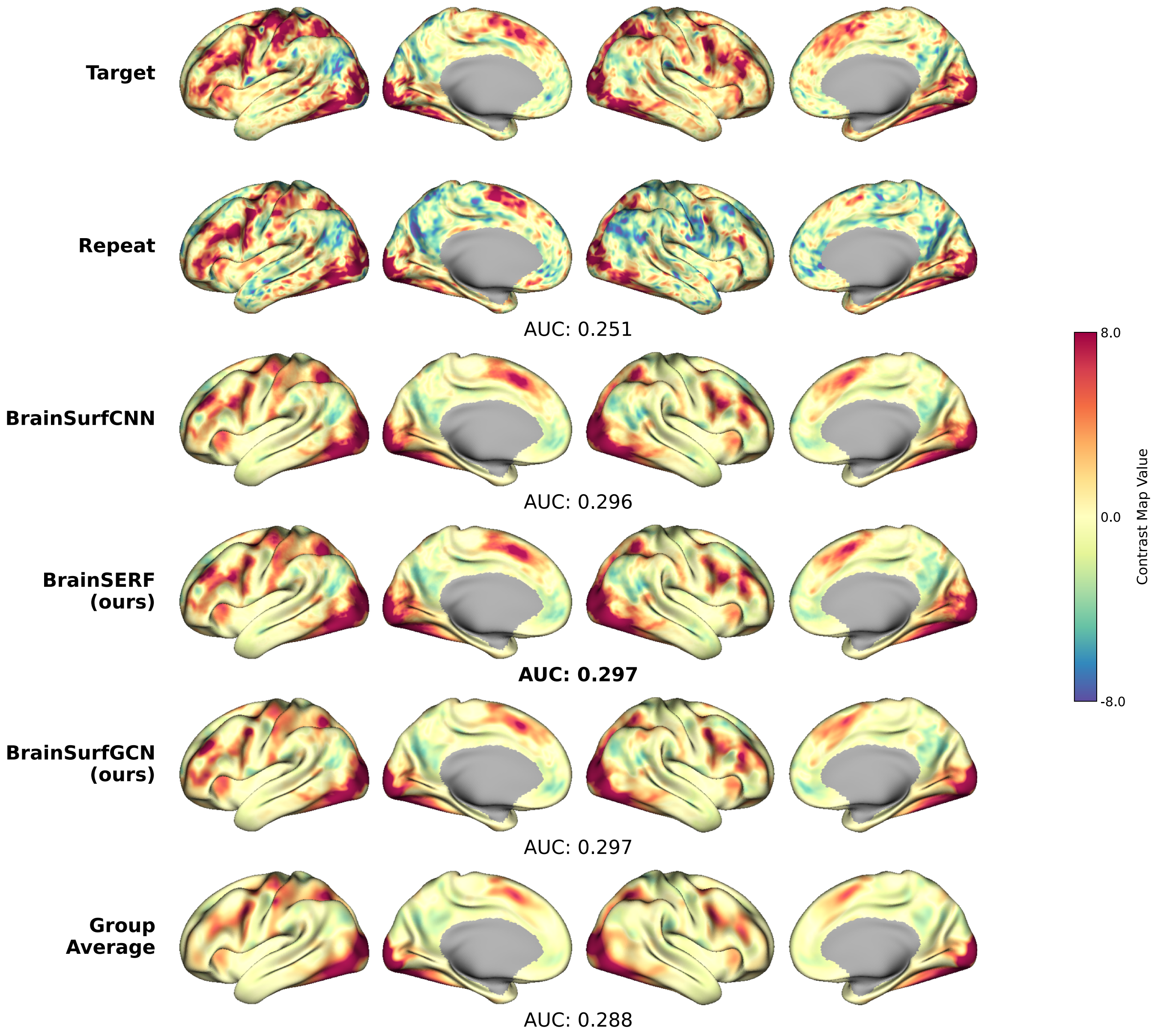


Figure S9: **Examples of unthresholded task activation for Working Memory: 0-Back.** We compare ground truth (target), repeat, model-predicted activation, and group average for an individual (subject 917255).

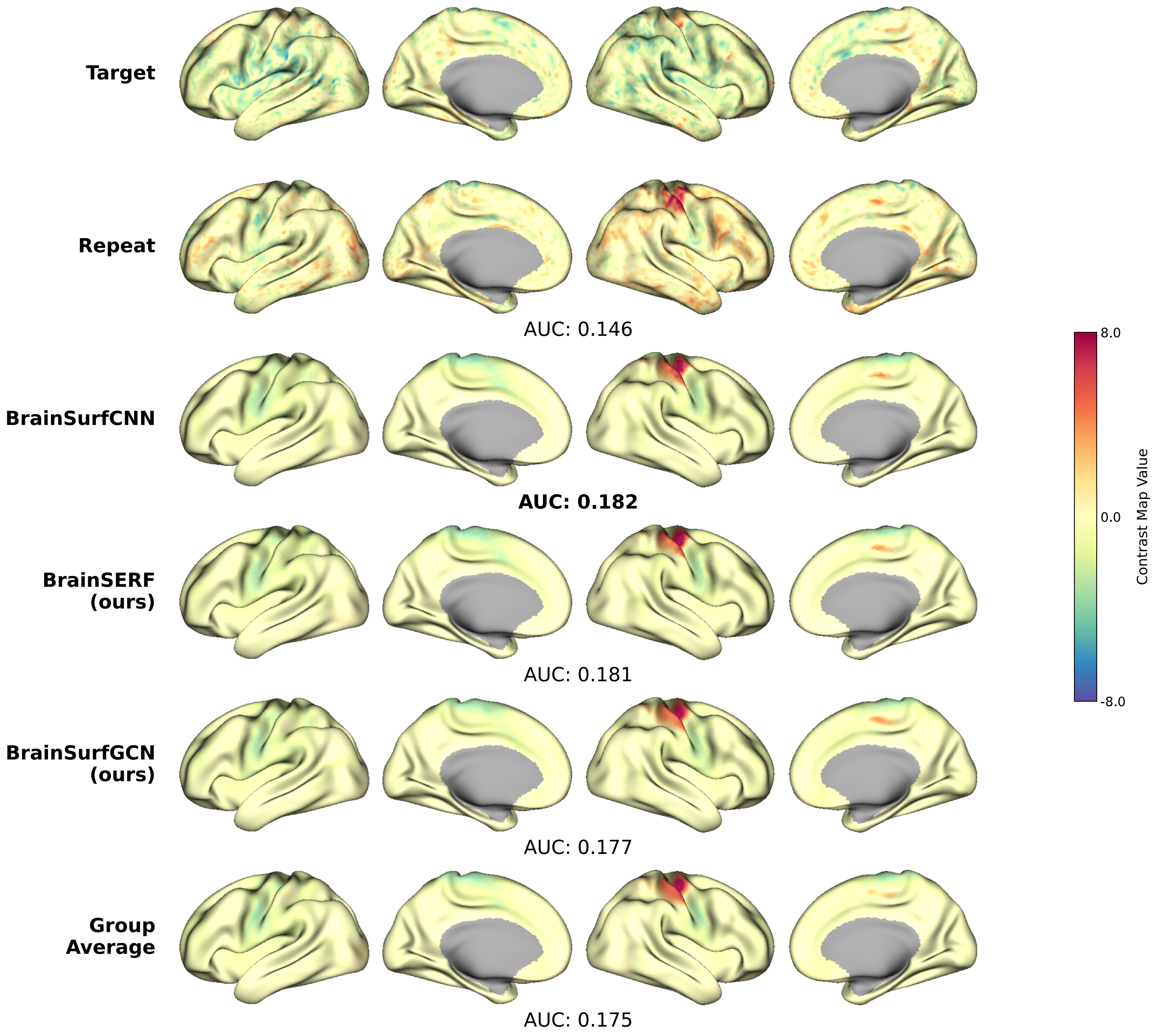


Figure S10: **Examples of unthresholded task activation for Motor: Left Hand-Average.** We compare ground truth (target), repeat, model-predicted activation, and group average for an individual (subject 917255).

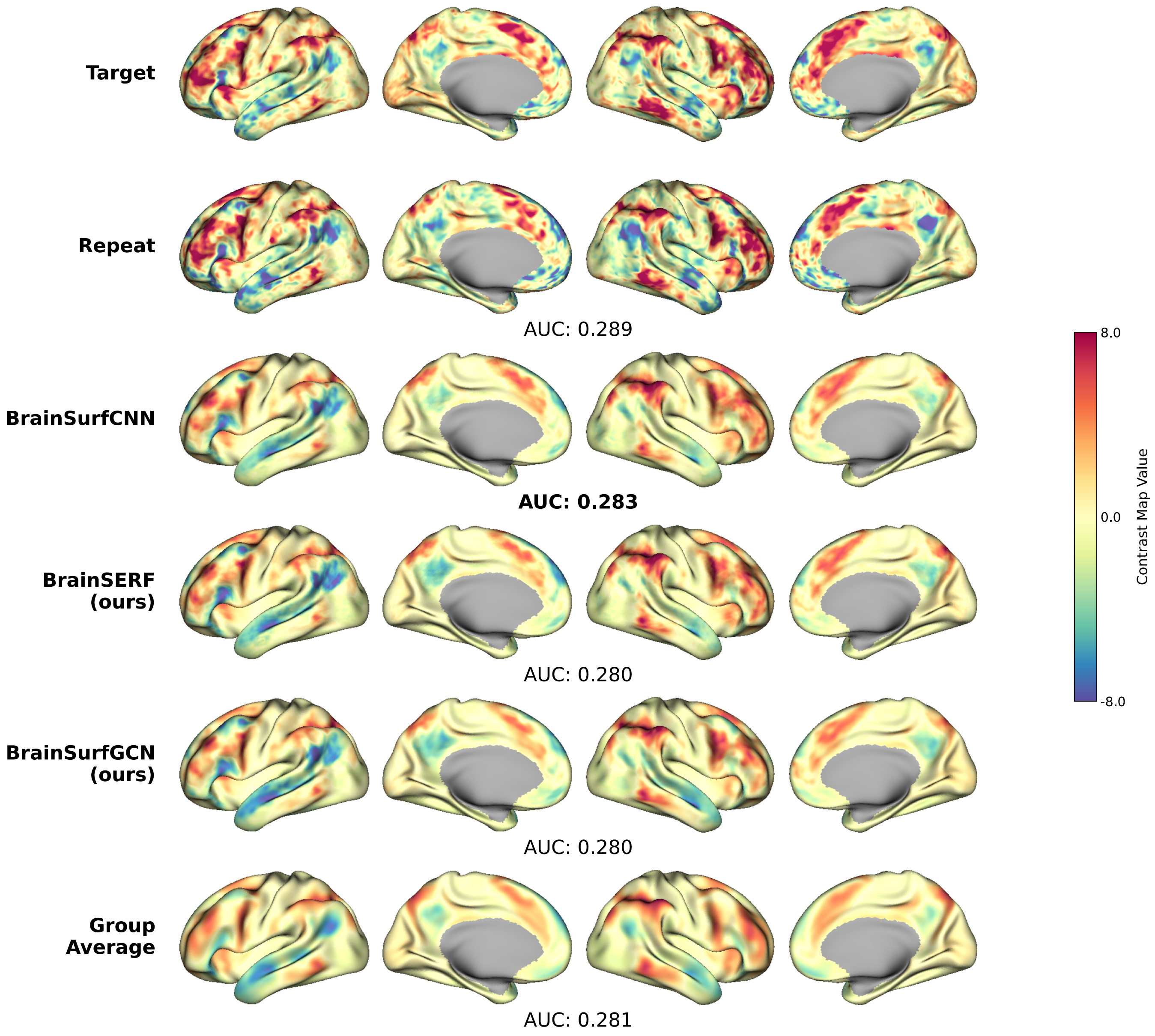


Figure S11: **Examples of unthresholded task activation for Language: Math-Story.** We compare ground truth (target), repeat, model-predicted activation, and group average for an individual (subject 103818).

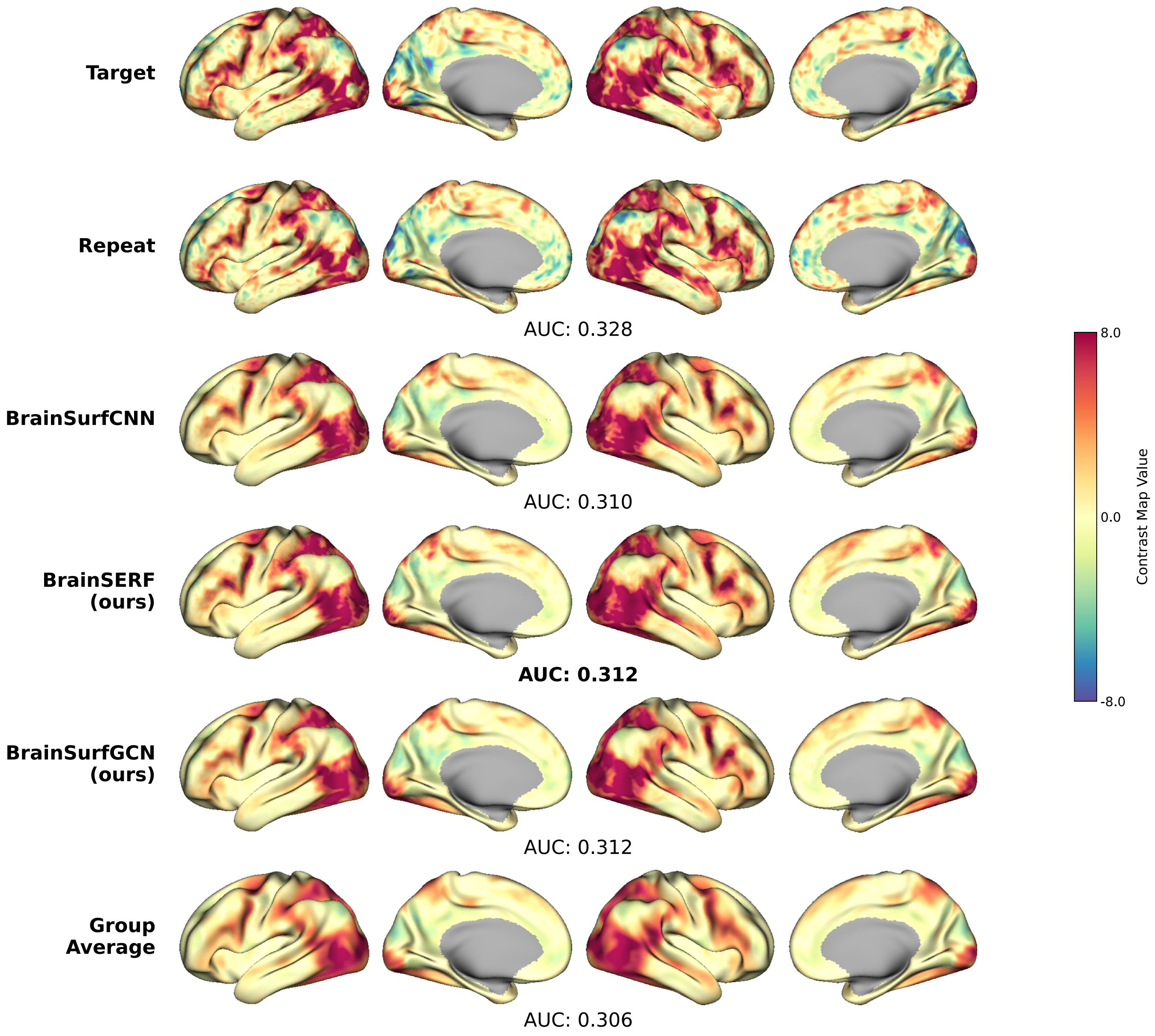


Figure S12: **Examples of unthresholded task activation for Social Cognition: Theory of Mind.** We compare ground truth (target), repeat, model-predicted activation, and group average for an individual (subject 103818).

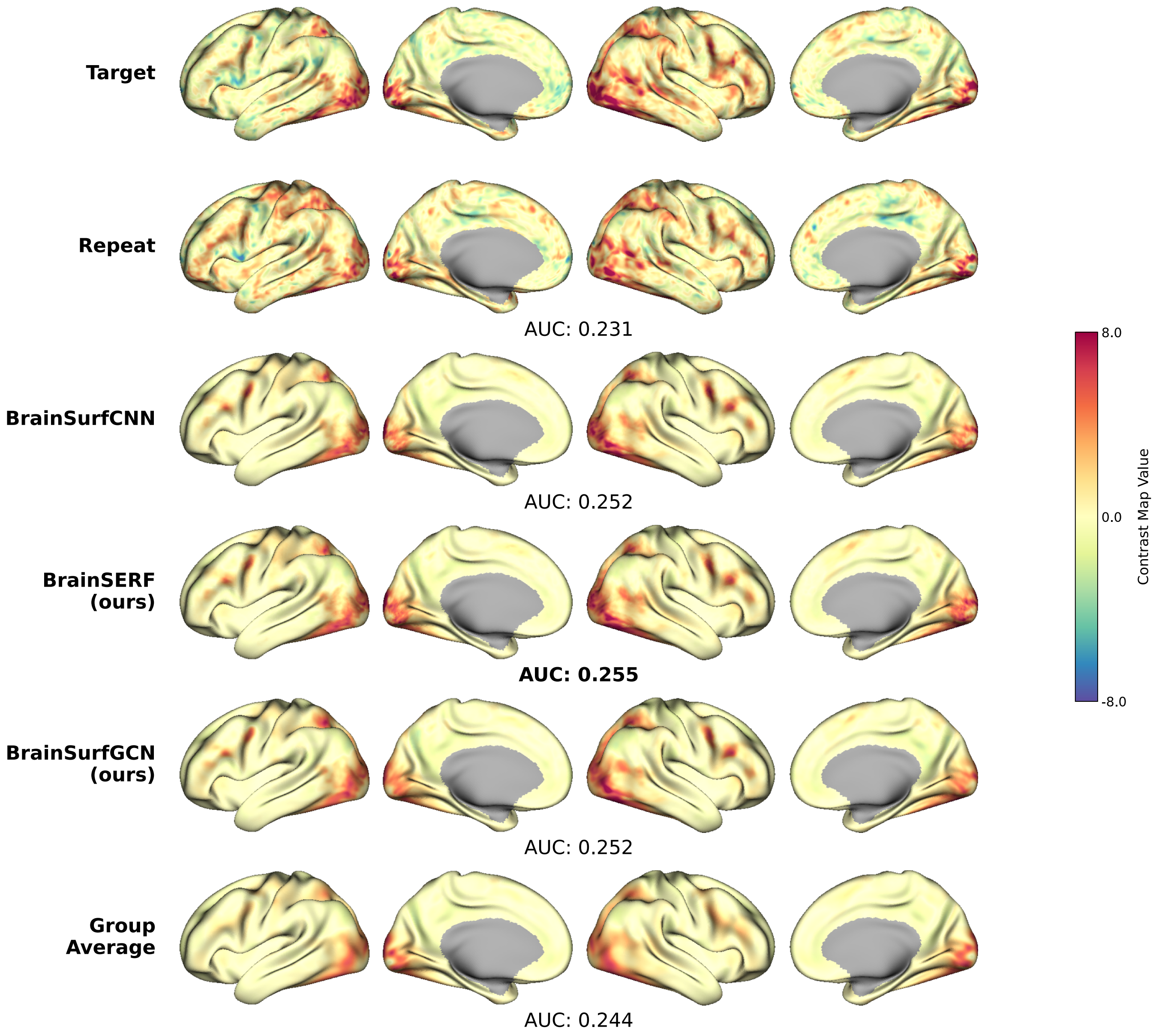


Figure S13: **Examples of unthresholded task activation for Emotion: Face.** We compare ground truth (target), repeat, model-predicted activation, and group average for an individual (subject 103818).

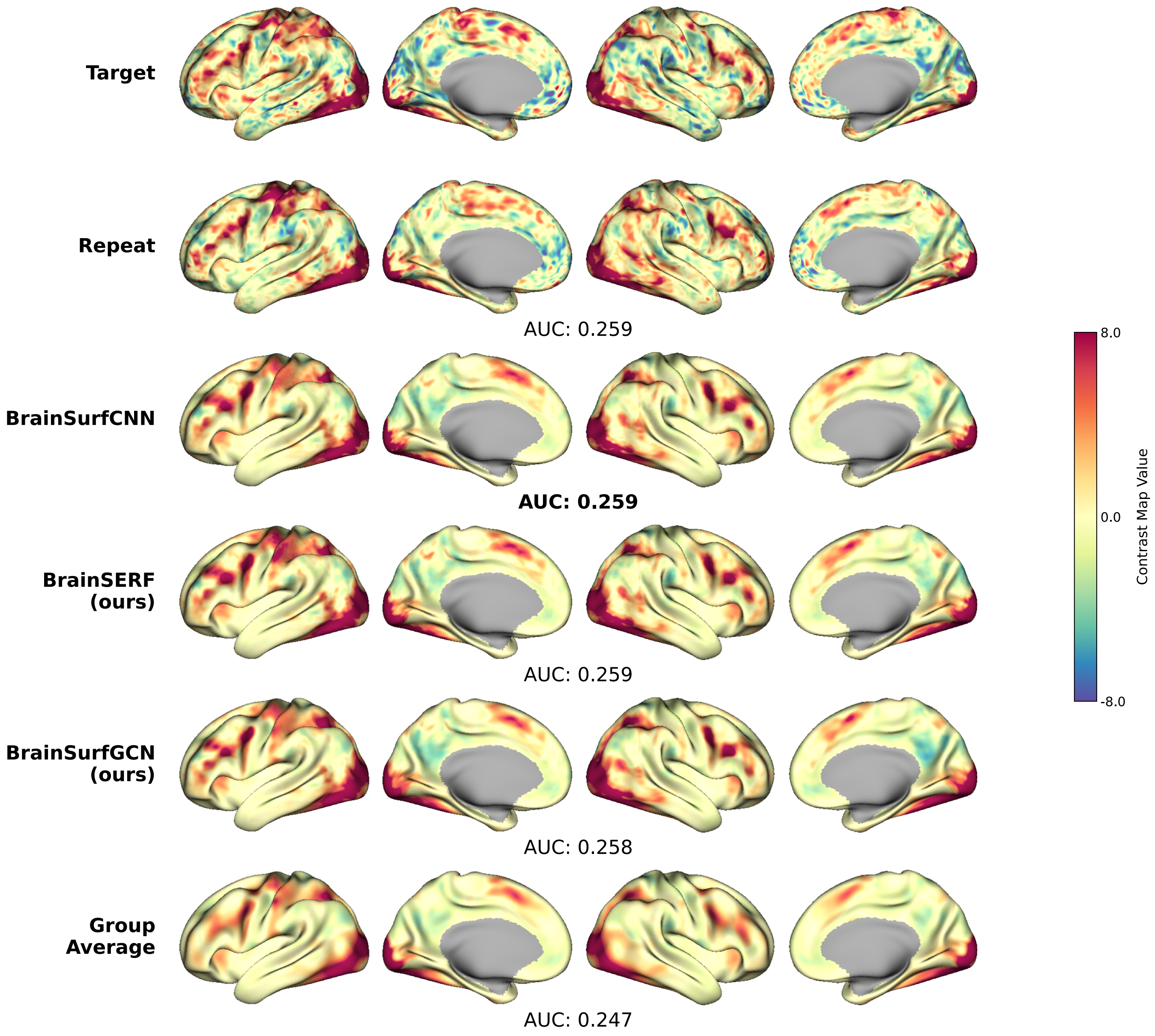


Figure S14: **Examples of unthresholded task activation for Working Memory: 0-Back.** We compare ground truth (target), repeat, model-predicted activation, and group average for an individual (subject 103818).

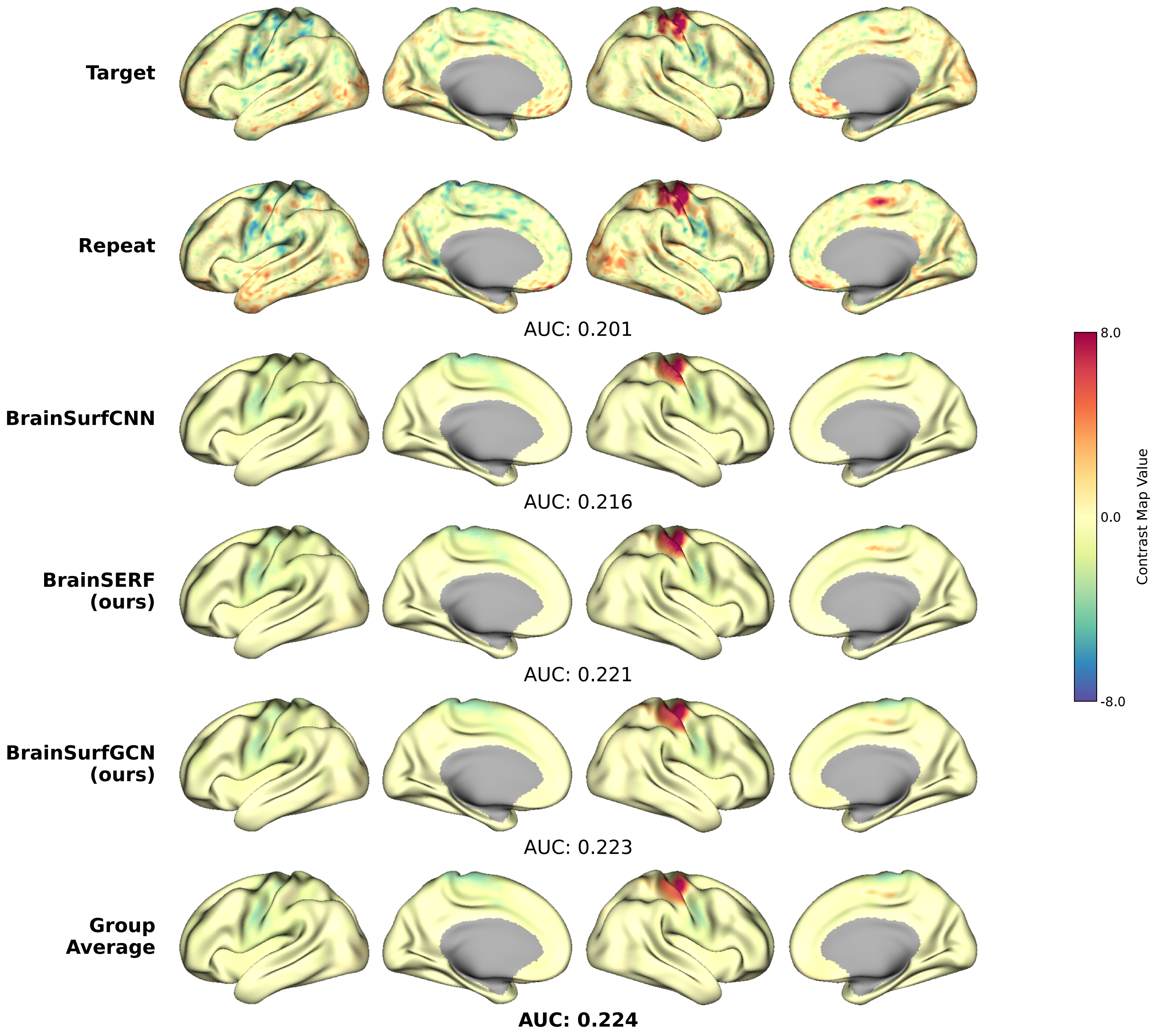


Figure S15: **Examples of unthresholded task activation for Motor: Left Hand-Average.** We compare ground truth (target), repeat, model-predicted activation, and group average for an individual (subject 103818).

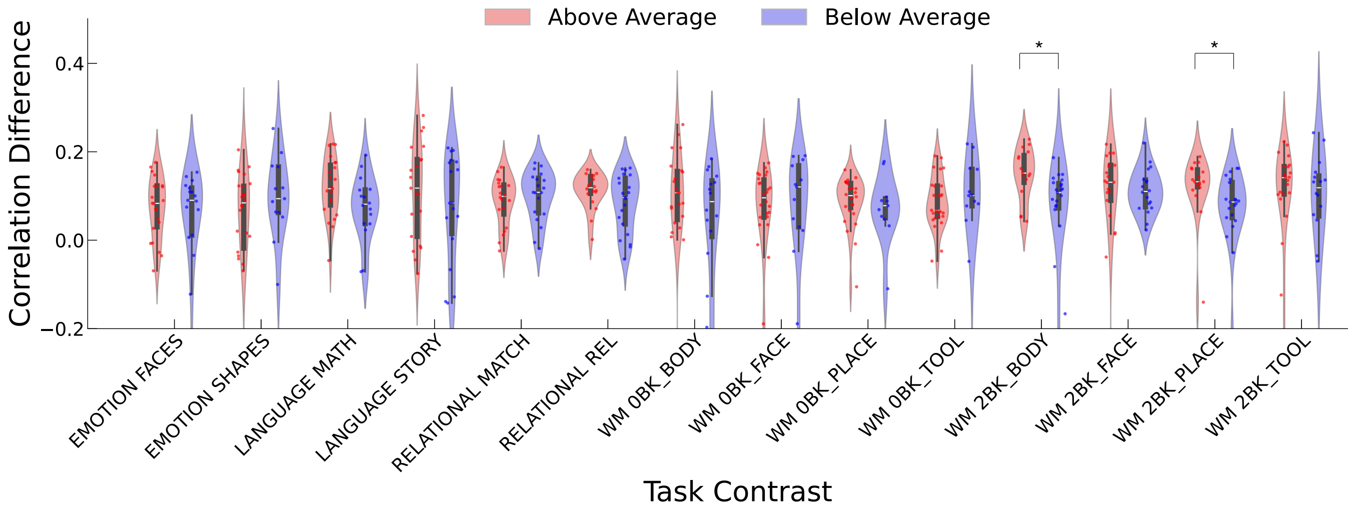


Figure S16: **Correlation difference for above- and below-average task accuracy groups across all task contrasts (BrainSurfGCN).** Violin plots show the distribution of correlation difference for subjects with above-average (red) and below-average (blue) behavioral performance across all task contrasts with an available accuracy metric. After Benjamini-Hochberg FDR correction, only Working Memory: 2-Back Body and Working Memory: 2-Back Place exhibited significant group differences.

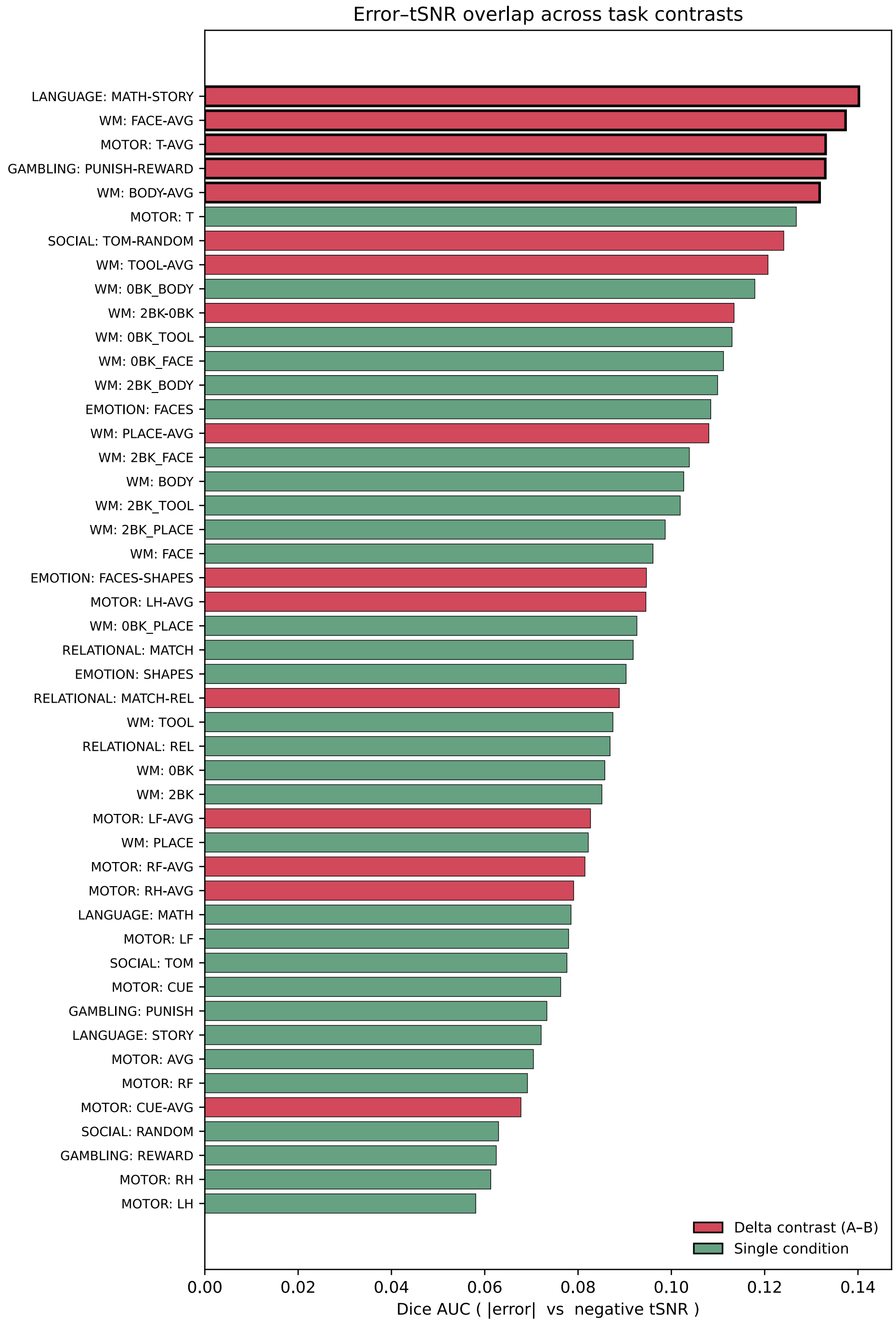


Figure S17: **Spatial overlap between rest-to-task prediction error and resting-state tSNR across all task contrasts.** Dice AUC between top-$t$ vertices of the absolute test-retest error map and the negative tSNR map (threshold sweep $t \in [0.05, 0.50]$), shown for all 47 HCP contrasts sorted in descending order. Delta contrasts (red) and single-condition contrasts (green); top-5 outlined in bold. Delta contrasts show significantly higher overlap (mean $0.108$ vs $0.088$; permutation $p = 0.001$, Cliff's $\delta = 0.48$).
